# A *Taphrina* strain infecting the model plant *Arabidopsis thaliana*

**DOI:** 10.1101/2021.09.16.460675

**Authors:** Margaretta Christita, Agate Auzane, Kai Wang, Timo Sipilä, Petri Auvinen, Lars Paulin, Jarkko Salojärvi, Kirk Overmyer

**Affiliations:** Organismal and Evolutionary Biology Research Program, Faculty of Biological and Environmental Sciences, and Viikki Plant Science Centre, University of Helsinki, Finland; Environment and Forestry Research and Development Institute of Manado, Indonesia; Finnish Institute of Molecular Medicine, P.O. Box 20, University of Helsinki, FI-00014 Helsinki, Finland; Institute of Biotechnology, University of Helsinki, Helsinki, Finland; Population Genomics and System Biology, School of Biological Sciences, Nanyang Technological University, 60 Nanyang Drive, SBS-01n-21, Singapore 637551, Singapore

## Abstract

Yeasts are important plant-associated organisms that can modulate host immunity to either promote or prevent disease. Mechanisms of plant-yeast interactions, specifically of yeast perception by the plant innate immune system, remain unknown. Progress has been hindered by the scarcity of yeast species associated with the model plant *Arabidopsis thaliana* (*Arabidopsis*). We have previously isolated *Taphrina* strain M11 from wild *Arabidopsis* in the field. *Taphrina* are poorly studied dimorphic yeast-like fungi that are plant pathogens, often producing plant hormones and causing tumour-like and leaf deformation symptoms on their hosts. Here we characterize the interaction of M11 with *Arabidopsis*. Infection of *Arabidopsis* with the birch pathogen *T. betulina*, used as a non-host control, shows early HR, enhanced ROS accumulation, and limitation of growth, demonstrating that *Arabidopsis* had immunity against non-adapted yeasts. M11 triggered limited cell death, an attenuated ROS response, and grew *in planta*, as well as subtle but clear leaf deformation symptoms, demonstrating it is pathogenic. Hormone responsive promoter-reporter analysis demonstrated activation of cytokinin signalling during infection. Mutant infection assays indicate jasmonate and ethylene were required for immunity against M11. Analysis of the *Taphrina* M11 genome was used to mine evidence for yeast specific PAMPs which may underlie host immune responses against yeast-like fungi.

## Introduction

Pathogens are specifically adapted to overcome specific host and non-host resistance mechanisms utilized by the host to limit their access or growth (Pieterse et al., 2012, Dodds and Rathjen, 2010, Cui et al., 2015; Lee et al., 2017). Plants possess a two-tiered innate-immune system (Jones and Dangl, 2006). The first level is termed basal or non-host resistance and consists of multiple mechanisms (Lee et al., 2017), including pattern recognition receptors (PRRs), which recognize conserved pathogen associated molecular patterns (PAMPs) to induced PAMP-triggered immunity (PTI) (Jones and Dangl, 2006). To overcome PTI, adapted pathogens have evolved effectors, which are toxins or short secreted proteins that interfere with plant immune signalling. To counter this plants deploy nucleotide-binding, leucine-rich repeat (NLR) immune receptors, which are activated by effector action leading to recruitment of immune signalling pathways, termed effector triggered immunity.

Non-pathogenic yeast species, such as *Saccharomyces cerevisiae* and *Schizosaccharomyces pombe* have been used as model system for genetics and molecular biology because of their easy genetics, rapid growth, and ease of cultivation. These model systems have been invaluable reference organisms in fungal genetics; for instance in definition of the unique components found in yeast cell walls (Perez et al., 2018). Human pathogenic yeasts such as *Candida albicans* have been important systems for understanding yeast-host interactions and have resulted in definition of the human PRRs responsible for monitoring the presence of pathogenic yeasts (Jouault et al., 2009). There are very few known phytopathogenic species among the true yeasts in subphylum Saccharomycotina, with a few examples in the genus *Eremothecium* (Wendland and Walther, 2011). However, there are a large number of phytopathogenic yeast-like fungi with dimorphic lifecycles, for instance in the Ascomycete subphylum Taphrinomycotina and the Basidiomycete subphylum Ustilaginomycotina (Begerow et al., 2017; Lachance and Walker, 2018).

Species belonging to the genus *Taphrina* (Ascomycota, Taphrinomycotina, Taphrinomycetes, Taphrinales, Taphrinaceae) are little studied pathogens of primarily woody plant species, although some *Taphrina* are pathogens of herbaceous host species; including for example, *Curcuma*, *Potentilla*, and some ferns (Mix, 1949; Ahmed and Kulkarni, 1968; Fonseca and Rodrigues, 2011; Rodrigues and Fonseca, 2003). *Protomyces* is the sister genus to *Taphrina*; members of this genus are pathogenic mostly on herbaceous hosts in the Umbelliferae and Compositae and have similar lifecycles and virulence strategies to the *Taphrina* (Wang et al., 2021). *Taphrina* are dimorphic, with dikaryotic infectious filamentous phase, which invades host tissues, and an easy to culture haploid yeast phase, which resides in the phyllosphere of the host between infection cycles. The *Taphrina* infection process has been shown to be dependent on cold and moist conditions during the infection window in spring while host buds opening (Giosuè et al., 2000; Rossi et al., 2006). Thus, infections do not necessarily occur on a yearly basis and it is held that *Taphrina* can survive as a yeast in the host phyllosphere indefinitely (Mix, 1949). Most *Taphrina* isolates have been isolated from their respective hosts, in a diseased state (Mix, 1949). Some *Taphrina* have been isolated in their yeast states from inert surfaces or plants without disease symptoms, suggesting that some *Taphrina* may be specialized in atypical lifestyles, such as endoliths or non-pathogenic phyllosphere residents (Selbmann et al., 2014; Inacio et al., 2004; Moore, 1998). Much of the previous work on members of the genus *Taphrina* is quite old; however, recent genome sequencing projects have opened this genus to modern molecular approaches (Cissé et al., 2013; Tsai et al., 2014; Wang et al., 2020).

Most *Taphrina* cause tumour-like symptoms on their hosts, as are typified by the witches’ brooms caused by the birch pathogen, *T. betulina* (Dingley, 2012; Bacigálová et al., 2005; Mix, 1949; Kern and Naef-Roth, 1975), or the leaf deformations caused by the peach pathogen, *T. deformans* (Mix, 1949; Fonseca & Rodrigues, 2011). *Taphrina* species are known for the ability in produce plant hormones auxin and cytokinin, but the role of these hormones in *Taphrina* pathogenicity has not been confirmed. Production of auxins and cytokinins by *T. betulina* was first reported in 1975 (Kern and Naef-Roth, 1975), while auxin production by *T. wiesneri* was first reporedt in 1966 (Matuyama and Misawa, 1961). The IAA biosynthesis pathways utilized in several *Taphrina* and *Protomyces* species have been addressed in several publications including genome studies (Cissé et al., 2013; Tsai et al, 2014; Wang et al., 2019). Additionally, secondary metabolite biosynthesis gene clusters have also been identified in the *T. deformans* genome (Cissé et al., 2013).

*Arabidopsis thaliana* (referred to here as *Arabidopsis*) is a widely used genetic model plant, which has accelerated definition of the plant immune system. Many studies have utilized the genetic model plant *Arabidopsis* to investigate the relationship of the plant innate immune system facing diverse microbes, including fungi, bacteria, oomycetes, but not yeasts. While PRRs and NLRs involved in immunity against other pathogen classes are well defined, the immune receptors detecting yeasts remain unknown. The investigation of yeast-plant interaction by utilizing *Arabidopsis* has been slowed by a lack of *Arabidopsis* associated yeasts. We have previously isolated *Taphrina* strains from wild *Arabidopsis* (Wang et al., 2016) opening the possibility to study *Arabidopsis* interactions with a pathogenic yeast. Previous studies have used *T. betulina* to study the non-host interaction with *Arabidopsis* (Gehrmann, 2013) and *Protomyces arabidopsidicola* to probe *Arabidopsis* immune activation by a phyllosphere resident non-pathogenic yeast (Wang et al., 2019; Wang et al., 2021).

*Arabidopsis* has not been previously known to be a host for *Taphrina*. In this study we describe *Taphrina* strain M11 isolated from wild *Arabidopsis* and compare it to known *Taphrina* species.. We describe *Arabidopsis* infection symptoms and potential interaction mechanisms with *Arabidopsis* immunity, such as alteration of plant hormone levels.

## Materials and Methods

### Plants and cultivation conditions

Wild type Col-0 accession and mutant seeds of *Arabidopsis* (*Arabidopsis thaliana*) were obtained from the Nottingham Arabidopsis stock centre (NASC; http://arabidopsis.info/). The mutant lines used where; *coi1-16 (coronatine insensitive1-16*), *cyp79 b2/b3 (cytochrome p450, family 79, subfamily b polypeptide 2 and 3 double mutant)*, *ein2 (ethylene insensitive2)*, *jar1 (jasmonate resistant1)*, *npr1 (non-expresser of pathogenesis-related genes1*), *pad 4 (phytoalexin deficient4)*, *sid2 (salicylic acid induction deficient2)*. All mutant alleles are in the Col-0 accession.

Standard plant cultivation conditions were as follows. For soil grown plants, seeds were sown on a well watered 1:1 mix of peat (Kekkilä; www.kekkila.fi) and vermiculite, stratified in the dark at 4°C for 72 hrs, then transferred to a growth chamber (Fitotron SGC120, Weiss Technik; www.weiss-technik.com) with 8/16 light hours of 150 uM M^-2^ illumination, at 23°C and constant humidity (ca. 60%). For sterile plant cultivation, seeds were sterilized with chlorine gas for 5h, sown on 0.5xMS 0.8 % agar, and stratified in the dark at 4°C for 72 hrs. One-week-old seedlings were transplanted into 6-well plates with 4 ml of 0.5xMS 0.8 % agar. The medium and roots were separated from the shoots using tight fitting polypropylene disks with a 4mm hole in the middle. Plants were grown in the Fitotron SGC120 growth chamber with 12/12 h light/dark cycle at ∼170 μmol m^-2^ s^-1^ light, +23/+18°C, and 65/75% relative humidity.

### Microbe strains and culture conditions

*Taphrina* strain M11 was previously isolated from the phyllosphere of wild growing *Arabidopsis* (Wang et al., 2016). Strain M11 has been deposited in the HAMBI - Helsinki Microbial Domain Biological Resource Centre - under the accession number HAMBI: H3698 and in the DSMZ - The German Collection of Microorganisms and Cell Cultures - under the accession number DSM 110146. All other *Taphrina* strains were obtained from the Portuguese yeast culture collection (PYCC; https://pycc.bio-aware.com/). *T. betulina* (strain PYCC 5889=CBS 119536) is not adapted to *Arabidopsis* and was used as a non-host response control. Two *T. tormentillae* strains (strain CBS 332.55=PYCC 5705 and strain PYCC 5727), are the strains most closely related to M11. *T. tormentillae* strain PYCC 5705/CBS 332.55 (formerly named *T. carnea)* was originally isolated from birch and thought to be a birch pathogen, but later shown to be conspecific with *T. tormentillae* (Fonseca and Rodrigues, 2011). All yeast strains were grown on 0.2 x potato dextrose agar (PDA) made with 15 g/l agar in potato dextrose broth (PDB; BD Biosciences; https://www.bd.com). *Pseudomonas syringae* pv. tomato strain DC3000 (*Pst* DC300) transgenically bearing the *AvrB* avirulence gene was cultured in NYGA media.

### Arabidopsis infections

For infections of soil grown plants seven-day-old *Taphrina* cells were collected using an inoculation loop, washed in 2ml 10 mM MgCl_2_ and resuspended in the same at OD = 0.3. Leaf halves from four-week-old soil-grown *Arabidopsis* were hand inoculated with yeast suspensions using a needleless syringe, then returned to standard growth conditions, covered at high humidity for the first 24 hours. Similarly prepared suspensions (OD = 0.1) from a one-day-old culture of *Pst* DC300 *AvrB* were used as a positive control for HR induction. Mock treatments with 10 mM MgCl_2_ were used as a negative control in all infection experiments. Experiments with *Arabidopsis* immune signaling mutants were photographed at 10 dpi to document symptoms, visually evaluated, and cell death was visualized by trypan blue staining.

For *Taphrina* growth tests on sterile plants freshly grown cells of all strains were harvested, washed, and suspended in 10mM MgCl_2_ with 0.04% wetting agent Silwet L77. Yeast suspensions (OD_600_ = 0.02) and control solution were applied onto 24***-***day***-***old plants of wild type Col-0 using sterile plastic spray bottles. To ensure uniform yeast distribution onto plants, all wells except one were covered with sterile aluminum foi**l** and the spray bottle was kept at a constant distance from plates. Half of the plants were harvested immediately and the rest at 9 dpi. Harvested plants were put in tared tubes with 1 ml of 10mM MgCl_2_, weighed, cooled on ice, and ground in a tissue homogenizer (*Precellys* 24; https://www.bertin-instruments.com) with 3 mm silica beads (2 × 30 s at 5000 rpm, with cooling on ice between runs). Dilutions of homogenized plant samples were plated on 0.2x PDA and *Taphrina* colonies were counted after 4 days. Additionally, the gnotobiotic status of the seedlings was periodically checked by plating ground uninfected leaf samples on LB media and 1xPDA.

### Histological staining

Biofilm formation was quantified using the crystal violet staining method as in (Wilson et al., 2017). Four-day-old cells of the indicated *Taphrina* strains were harvested from 0.2x PDA media, washed, and suspended at OD_600_ = 0.02. 200 μl of yeast suspension were added to 800 μl of 0.2x PDB (final OD_600_ = 0.004) and grown in polystyrene 48-well plates (CELLSTAR® Cell Culture Multiwell Plates, TC treated, Greiner Bio-One) without shaking. To prevent contamination, each yeast species was separated by empty wells containing only media. After 8 and 16 days, wells were rinsed to remove loosely adherent cells, stained for 15 min with 1% crystal violet solution, and photographed. Subsequently, biofilm-bound dye was dissolved in 100% ethanol and quantified spectrophotometrically (λ600).

Crystal violet staining was used, as in Valadon et al (1962), to visualize M11 cells and biofilms in and on infected leaves; briefly, 0.5% crystal violet staining solution was prepared in aqueous 20% methanol. Leaves of wild type *Arabidopsis* that had been infiltrated with *Taphrina* strain M11 at 3-7 dpi were cleared in 90% acetone, placed on a slide, and stained with a drop of the staining solution for 5-10 sec at room temperature, destained with ddH_2_O as required, and mounted in 60% glycerol. Samples were observed under a Leica compound microscope (MZ 2500; https://www.leica-microsystems.com)with a magnification of 200x-400x. Trypan blue staining was used to visualize hypersensitive response-like cell death in infected soil grown plants. Whole leaves were stained by boiling for ca. 1 minute in a 1:1 dilution of trypan blue staining solution (0:05%) in lactophenol (1:1:1, glycerol:85% lactic acid: phenol) in 95% ethanol. Samples were cleared at room temperature in chloral hydrate (2.5g chloral hydrate per 1ml ddH_2_0) with several changes of destaining solution until clear.

DAB (3,3’-Diaminobenzidine; Sigma; www.sigmaaldrich.com) staining was used to visualize the *in planta* accumulation of H_2_O_2_ (Jambunathan, 2010). DAB stain solution (0.1% w/v) was prepared fresh and protected from light. Four-week-old *Arabidopsis* were hand inoculated and stained for 2 hrs in a closed container in the dark at high humidity. DAB solution was infiltrated into the leaves by vacuum infiltration. The staining reaction was stopped by immersing samples in clearing solution (3:1 solution of 95% ethanol in lactophenol).

β-Glucuronidase (GUS) staining was used to investigate activation of host plant auxin and cytokinin transcriptional responses during infection using transgenic Col-0 *Arabidopsis* with the following promoter-reporter systems; the auxin-responsive DR5 promoter or the cytokinin responsive TCS promoted fused to the GUS reporter; denoted as DR5::GUS and TCS::GUS, respectively. Four-week-old *Arabidopsis* were infected by hand infiltration. Positive controls for DR5::GUS were treated with 2, 5, and 10 μM indole acetic acid (IAA), for TCS::GUS controls were 2, 5, and 10 μM 6-benzylaminopurine (BAP), all negative controls were mock infected with MgCl_2_+ silwet. GUS staining solution contained 1 mM 5-bromo-4-chloro-3-indolyl b-D-glucuronide dissolved in methanol, 5 mM potassium ferricyanide and 5 mM potassium ferrocyanide in 50 mM sodium phosphate buffer and adjusted to pH 7.2. For histochemical staining, seedlings were fixed with ice-cold 90% acetone for 1 h, washed two times with ice-cold wash solution (50 mM sodium phosphate buffer, pH 7.2), 30 min for each wash. Seedlings were vacuum infiltrated for 5 minutes and kept at room temperature in GUS staining solution. Stained seedlings were washed two times with absolute ethanol, then cleared and stored in 70% ethanol.

### Leaf symptom assays

Leaf symptom assays included quantification of leaf curling and leaf bending. To investigate leaf curling, Col-0 leaves were infected with strain M11, and *T. betulina* at 14 dpi were transversely cross sectioned half way between the leaf base and tip by hand using razor blade and photographed. Curling index was measured from photos of leaf sections using Image J (https://imagej.nih.gov/ij/), as in Booker et al.(2004) , and as illustrated in Figure S1.

To quantify leaf bending, M11 and *T. betulina* infected Col-0 leaves were photographed at 14 dpi then leaf bending was measured using Image J. Leaf bending was quantified by measuring the angle between the base of the petiole and the leaf tip, as illustrated in Figure S1, were a greater the angle indicates a higher level of leaf bending.

### Genome sequencing, assembly, and analysis

DNA extraction, genome sequencing, and assembly were performed as previously described in Wang et al. (2019). In short, chromosomal DNA was extracted according to the protocol of Hoffman (1997). DNA quantity and quality was assessed using qubit fluorometer (Thermo Fisher Scientific, USA) and Nanodrop ND-1000 (Thermo Scientific, USA). Following the Paired-End Sample Preparation Guide (Illumina) DNA libraries were prepared for sequencing with MiSeq System (Illumina, California USA). Sequencing was performed at the DNA Sequencing and Genomics Laboratory, Institute of Biotechnology, University of Helsinki. Resulting reads were assembled with SPAdes v. 3.1.1 pipeline (Bankevich et al., 2012). Assembly quality was determined using the QUAST tool version 5.0 (http://quast.sourceforge.net/).

Genome annotation was performed using Augustus version 2.5.5 (Stanke et al., 2008) which was trained on RNA sequencing results from *Taphrina betulina* genome sequencing project (Bioproject: PRJNA188318). For further details on the automated annotation pipeline see Wang et al. (2019).

To analyse the distribution of orthologous proteins in *Taphrina* M11 and selected members of the *Taphrinomycotina* the OrthoVenn2 platform (Xu et al., 2019) was used with an E-value cut-off E < 0.01. Proteomes from the following whole genome sequencing projects were included: *T. deformans* JCM22205, BAVV01; *T. wiesneri* JCM22204, BAVU01; *T. populina* JCM22190, BAVX01; *T. flavorubra* JCM22207, BAVW01; *S. pombe* 972h-, ASM294v2. Proteomes of *P. arabidopsidicola* strain C29, QXMI01; *P. lactucaedebilis* YB-4353, QXDS01; *P. gravidus* Y-17093, QXDP01; *P. macrosporus* Y-12879, QXDT01 were from (Wang et al., 2019).

Conserved domains in annotated proteins of M11 were identified using HMMER software versions v3.2.1 and v3.3 by searching against Pfam database with E < 1e-30 cut-off (Finn et al., 2016).

To identify candidate effector-like proteins, open reading frames (ORFs) in the size range 80-333 amino acids (aa) were selected for screening with SignalP 4.1 tool for the presence of a secretion signal (Petersen et al., 2011), and defined as short secreted proteins (SSPs). SSP sequence secretion signals were trimmed and mature SPP peptides with ≥4 cysteine residues were categorized as cysteine-rich SSPs (CSSPs).

Orthologs for enzymes of interest were identified using BLASTp tool to search the M11 and other *Taphrina* genomes with the query sequences provided in the supplemental files. The identity of proteins was further confirmed by performing BLASTp search against Swiss-Prot database (default parameters). For the alignment of chitin synthases Clustal Omega multiple sequence alignment program was used (Madeira et al., 2019).

## Results

### Taphrina M11 growth on Arabidopsis

We have previously isolated a collection of yeasts from the phyllosphere of wild growing *Arabidopsis* in the field (Wang et al., 2016). These yeast species were screened for the ability to cause disease on *Arabidopsis*, including OTU3, which had two *Taphrina* strains, M11 and M12 with ITS sequences (LT602860) that are identical to each other and most closely related (99% similarity) to *T. tormentillae*, which is pathogenic on members of the herbaceous host plant genus the *Potentilla* (Wang et al., 2016; Fonseca & Rodrigues, 2011). Hand infiltration of *Arabidopsis* leaves with *Taphrina* strain M11 cell suspensions resulted in disease symptoms in *Arabidopsis* (Figure 1A), including leaf deformations and chlorosis. Based on this result, *Taphrina* strain M11 was targeted for genome sequencing and further characterization of its interaction with *Arabidopsis*.

**Figure 1.**
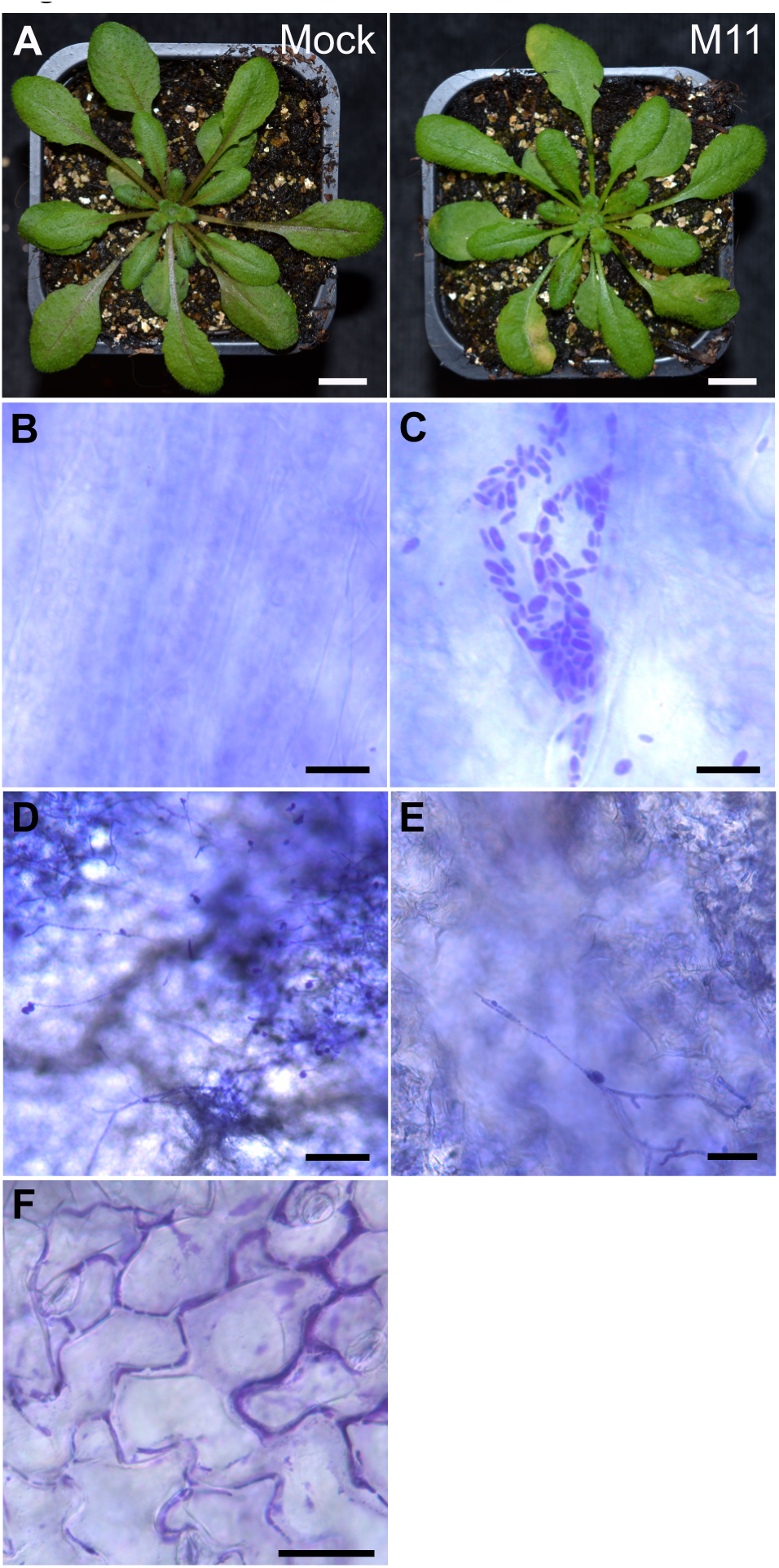
Host symptoms and *Taphrina* cell morphology in infected *Arabidopsis.* **(A)** Four-week-old control and infected *Arabidopsis* were photographed at seven days post infection (dpi). *Taphrina* strain M11 infiltration (M11), mock infiltration using MgCl_2_ (mock). Scale bars = 1 cm. **(B-F)** Visualization of the morphology of *Taphrina* strain M11 cells in the leaf of *Arabidopsis* after hand infiltration using a needless syringe with four-week-old *Arabidopsis*. Observations were made at one dpi and show a control mock infected leaf (Scale bar = 10 µm) **(B)**, and yeast cells (Scale bar = 10 µm). **(C)**, Hyphal cells were observed at three dpi (Scale bar = 5 µm). **(D)**, close up of hyphal cells (Scale bar = 2 µm). **(E)**, and biofilm formation observed on the leaf surface at three dpi, as shown, or later (Scale bar = 30 µm) **(F)**.

As *Taphrina* species have dimorphic lifecycles, we probed the morphology of strain M11 cells in *Arabidopsis* leaves using microscopic observation of crystal violet stained infected tissues. Comparison of uninfected control to infected leaf tissue revealed clusters of M11 yeast cells at 24 hpi (Figure 1B-C). In some tissues at slightly later time points (3 dpi) the growth of infectious hyphae was detected (Figure 1 D-E). The stain used here can also visualize biofilms (Wilson et al., 2017). In some infected *Arabidopsis* leaves, dark staining cell aggregates with the appearance of a biofilm were observed (Figure 1F).

M11 growth on *Arabidopsis* was then quantified from spray infections of axenic 24-day-old plants (Figure 2A) compared to the two closely related *T. tormentillae* strains (CBS 332.55 and PYCC 5727) and non-host response control, *T. betulina*. Strain M11 grew to the highest levels, closely followed by the two other *T. tormentillae* strains, which both grew to similar levels, while *T. betulina* showed little to no growth.

**Figure 2.**
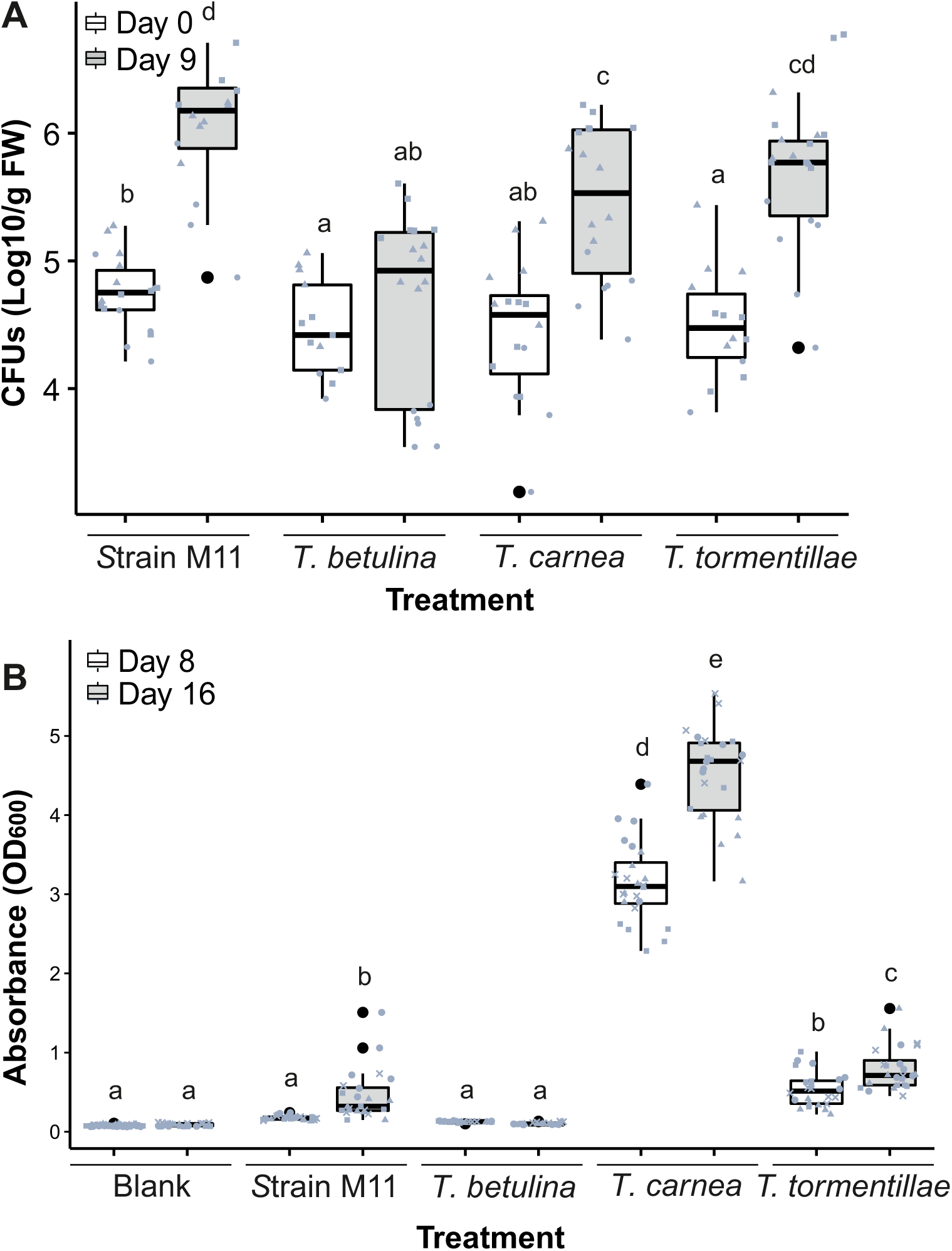
*Taphrina* growth and biofilm formation *in vitro.* *Taphrina* growth in the phyllosphere of *in vitro Arabidopsis* plants **(A)**. *Taphrina* yeast suspensions were sprayed onto the surface of 24-day-old, sterile wild type Col-0 accession *Arabidopsis* plants. Strains used were M11 *Taphrina*, the birch pathogen *T. betulina* used here as a non-host response control, and two strains of *T. tormentillae* - PYCC 5727/CBS339.55 (*T. tormentillae*) and PYCC 5705/CBC332.55 (*T. carnea*) which was originally assigned the name *T. carnea* but later shown to be a strain of *T. tormentillae*).Yeast growth was quantified through cultivation based technique immediately after spraying and at 9 dpi (days post infection). Combined data are presented from three independent experiments (n=6) for each experiment, data points from each experiment are represented with a different shape - circle, triangle, or square. Letters above boxplots represent significance groups from a Tukey’s test performed on linear mixed model computed in R with biological repeats as a random effect. Means α ≥ 0.05 share a common letter. **(B)** Biofilm formation by *Taphrina* species using the same strains as described above. The presence of adherent, biofilm-forming cells were monitored by spectrophotometric quantification of released crystal violet stain after treatment with a with 1% solution and ethanol extraction. Pooled data from four independent biological repeats is presented (n=6 for each biologcal repeat, total n=24), data points from each experiment are plotted with a different shape. Letters above boxplots represent significance groups as described above.

To further explore the possibility of biofilm formation by strain M11 the ability of *Taphrina* species to form biofilms was quantitatively assayed *in vitro. T. tormentillae* strain CBS 332.55 demonstrated strong ability to form biofilms on polystyrene surfaces (Figure 2B). In comparison, *T. tormentillae* strain PYCC 5727 showed medium biofilm formation on polystyrene and M11 started forming small, adherent biofilm-like patches only after 16 days of growth. The non-host control, *T. betulina,* was not able to form biofilms under the tested conditions.

### Response to infection

As shown in Figure 3A-D, the activation of cell death upon *Taphrina* infection was monitored visually (left) and with trypan blue staining at 48 hpi (right). Treatment of *Arabidopsis* leaves with M11 resulted in chlorosis and leaf deformation symptoms that were associated with a very low level of cell death (Figure 3A) compared to mock infected control leaves (Figure 3D). Activation of hypersensitive response (HR)-like cell death was observed in leaves challenged with the non-adapted *T. betulina* (Figure 3B); however, this was less than in leaves treated with avirulent *Pst* strain DC3000 transgenically bearing the *AvrB* avirulence gene, which was used as a control for a strong HR (Figure 3C).

**Figure 3.**
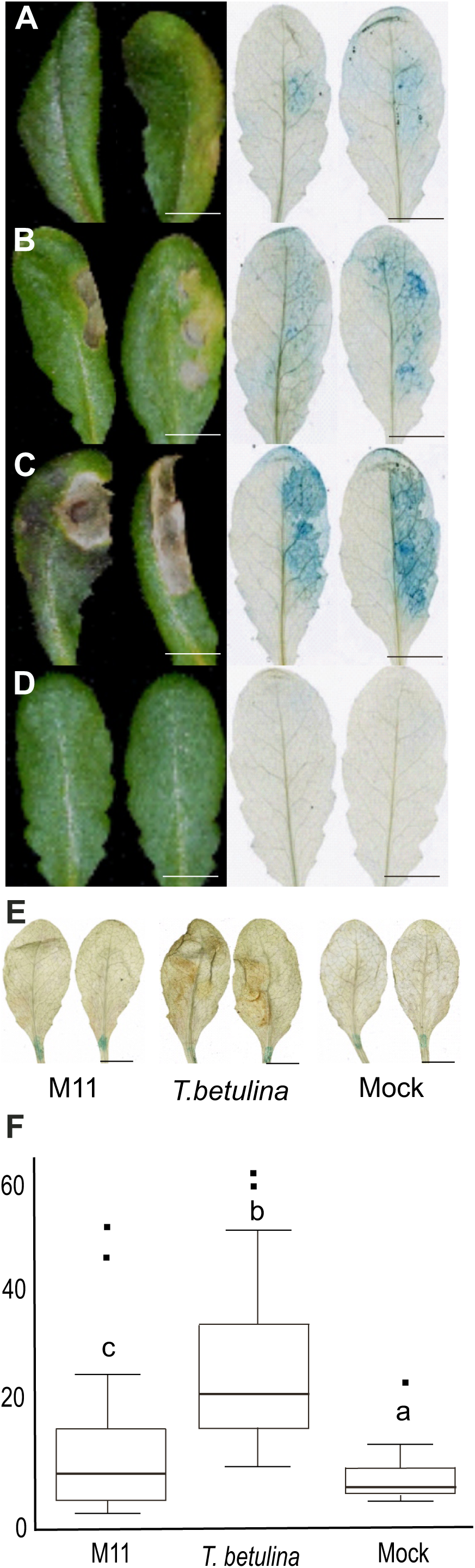
Cell death and ROS accumulation in infected *Arabidopsis.* Various cell suspensions were hand infiltrated using a needless syringe into leaf halves of four-week-old soil grown *Arabidopsis*. At 48 hours post infection (hpi), infected leaves were photographed (left) and trypan blue histological staining was used to visualize cell death (right). Leaves were infected with: **(A)** *Taphrina* strain M11, which was isolated from wild *Arabidopsis* **(B)** *T. betulina*, a birch pathogen used here as a control for a non-host response. **(C)** Avirulent *Pseudomonas syringae* pv. tomato DC3000 transgenically expressing *AvrB*, used as a positive control for the hypersensitive response **(D)** Mock (10 mM MgCl_2_). **(E-F)** ROS accumulation during infection of *Arabidopsis* with *Taphrina* strain M11 or *T. betulina* was monitored at 48 hpi by histologically visualizing H_2_O_2_ accumulation with 3,3’-diaminobenzidine (DAB) staining. Leaf halves of four-week-old soil grown *Arabidopsis* were hand infiltrated with a needless syringe. Stained and cleared leaves were photographed **(E)** and stain was quantified digitally by counting brown pixels in ImageJ **(F)**. One repeat representative of three independent biological repeats is shown. Letters above boxplots represent significance groups determined with ANOVA and Tukey HSD post-hoc test.

To monitor the accumulation of H_2_O_2_ caused by challenge with *Taphrina*, DAB staining was performed; stained and cleared leaves were photographed (Figure 3E) and brown colored pixels were quantified using ImageJ (Figure 3F). Compared to the mock treatment there was a small but significant increase in DAB staining in *Arabidopsis* infected with M11. In *T. betulina* inoculated *Arabidopsis* accumulation of DAB stain was significantly (p=2.106e-10) higher and covered a wider leaf area than in M11 infected plants (Figure 3E-F).

At later time points, *Arabidopsis* leaves infected with strain M11, also exhibited additional leaf symptoms, more subtle, but reminiscent of the leaf deformations caused by other *Taphrina* species; these late responses presented in the form of leaf curing and leaf bending (Figure 5). A leaf curling response was quantified using a curvature index as used previously (Booker et al., 2004), which is the ratio of the leaf width to the total leaf width (Figure S1A), where a smaller curling index value indicates a higher degree of leaf curvature (epinasty). Plants infected with the M11 strain exhibited visibly enhanced leaf curling compared to mock treated controls (Figure 4A), and quantified as a curling index 50%, while mock was 75% (Figure 4B). This phenotype was specific to strain M11, as infection with the non-host control, *T. betulina*, was indistinguishable from control both visually and quantitatively, with a curling index of 75% (Figure 4 A-B).

**Figure 4.**
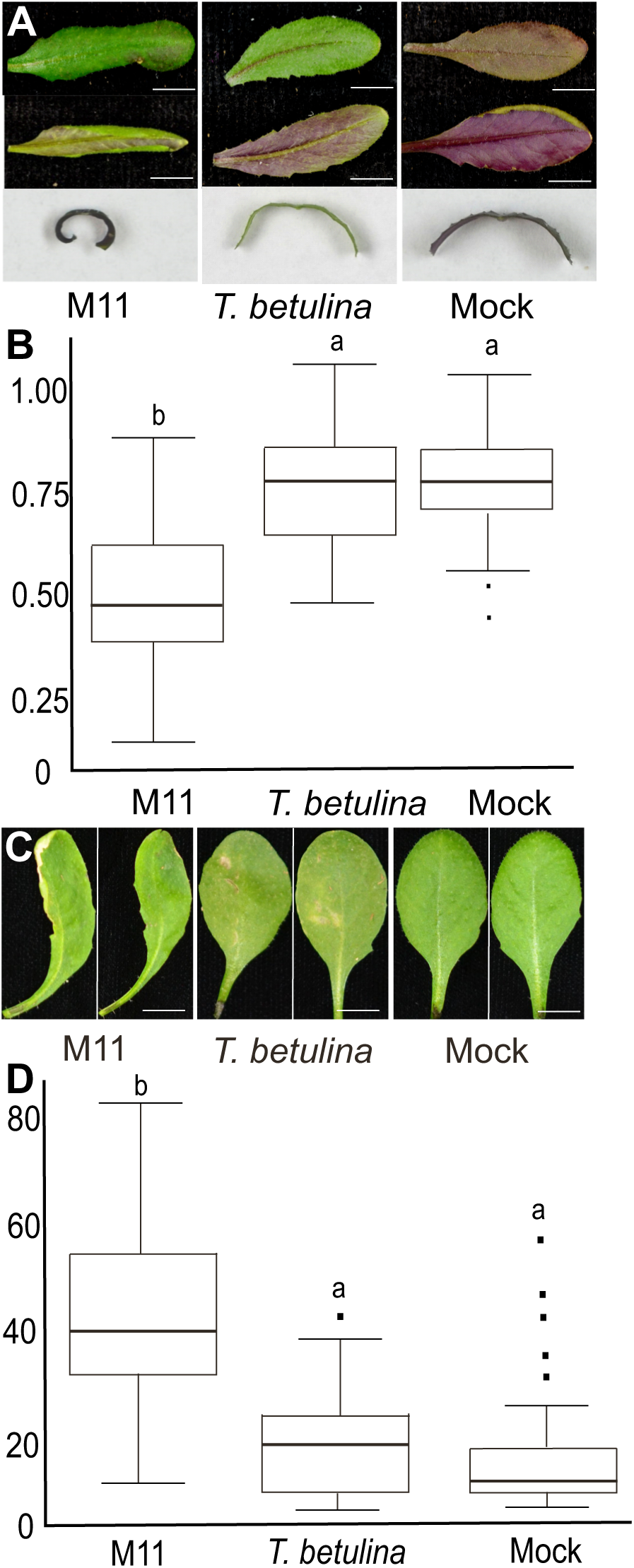
Leaf deformation phenotypes. Subtle leaf deformation phenotypes observed in response to infection with *Taphrina* species. Leaves of four-week-old soil grown Col-0 accession *Arabidopsis* were hand infiltrated with using a needless syringe delivering cell suspensions of M11, *T. betulina,* or mock treatment with10mM MgCl_2_. Observations were made at 14 dpi **(A)** Leaf curling was seen in primary infected leaves and in new leaves that developed after infection. Leaves were photographed on the adaxial side (top), abaxial side (middle) and hand sectioned at their mid-point. The ca. 1 mm hand section was placed on its side and photographed to reveal its curvature (bottom). Scale bars = 0.5 cm. **(B)** Leaf curing was quantitatively measured from the hand section photographs with ImageJ and the leaf curling index was calculated as the ratio of the leaf with and the total width (Figure S1A; (Booker et al., 2004), which results in lower scores for leaves with greater curvature. Box plots depict pooled results from three independent biological repeats (n= 5 per biological repeat, total n=15). Letters above the box plots indicate significance groups calculated with ANOVA and Tukey HSD post-hoc test. **(C)** Leaf bending phenotypes were documented in photographs of primary infected leaves. **(D)** Leaf bending phenotypes were quantitatively measured from photographs with ImageJ and the leaf curling index was calculated as the angle between a line defined by the petiole and a second line defined by the leaf mid-point and tip, as shown (Figure S1B). Box plots depict pooled results from three independent biological repeats (n=5 per biological repeat, total n=15). Letters above the box plots indicate significance groups calculated with ANOVA and Tukey HSD post-hoc test.

**Figure 5.**
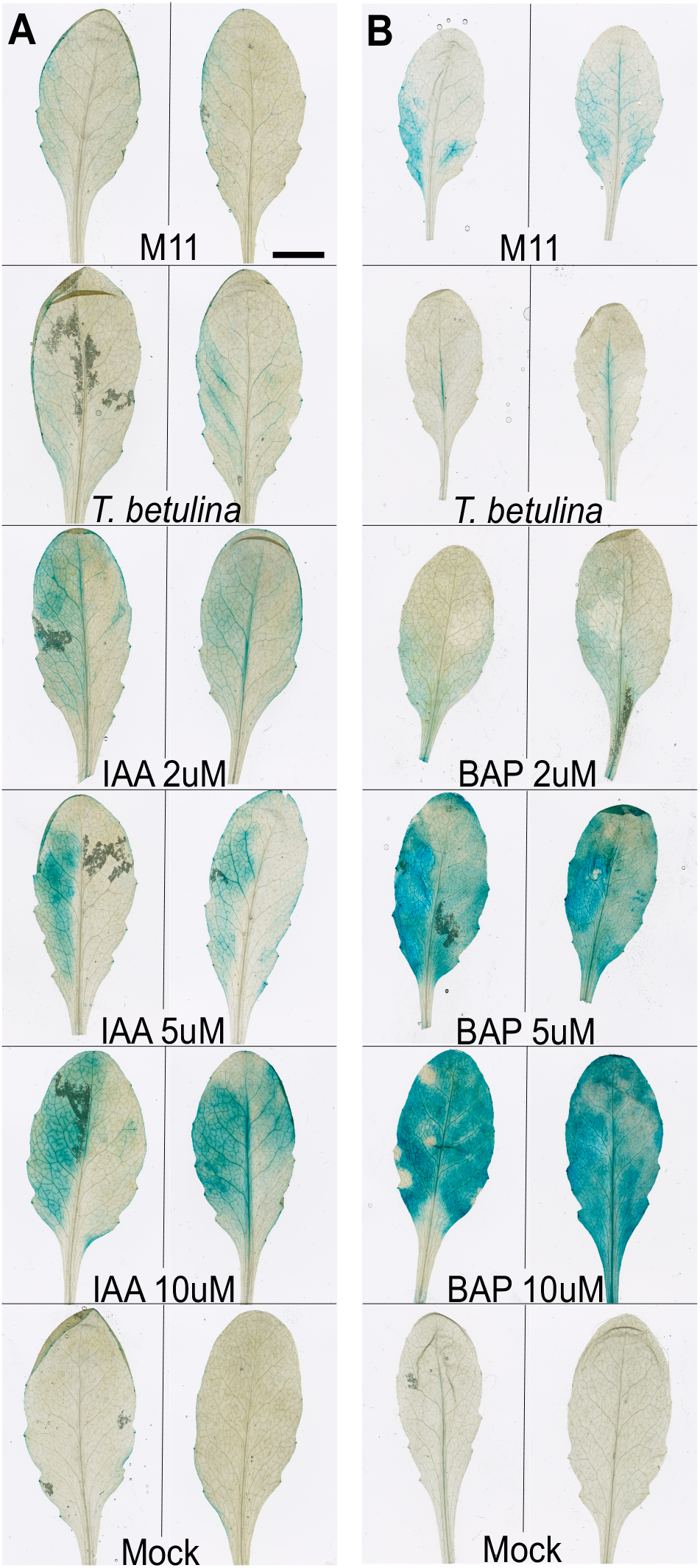
Activation of host hormone responses during *Taphrina* infections. Activation of the *Arabidopsis* auxin and cytokinin transcriptional responses in response to infection with *Taphrina* strain M11 or *T. betulina*. Infections of four-week-old soil grown *Arabidopsis was* by hand infiltration of *Taphrina* cell suspensions using a needless syringe. Left hand leave halves were infiltrated and histologically stained to visualize GUS activity at 48 hpi. **(A)** Activation of the plant auxin response to various treatments, as indicated, using plants transgenically bearing the auxin responsive pDR5::GUS promoter-reporter construct. As a positive control, treatments with the auxin, indole acetic acid (IAA), were used at the concentrations, 2, 5, and 10 uM and the negative control was mock infected with 10mM MgCl_2_. **(B)** Activation of the plant cytokinin response to various treatments, as indicated, using accession Col-0 plants transgenically bearing the cytokinin responsive pTCS::GUS promoter-reporter construct. As a positive control, treatments with the cytokinin, benzylaminopurine (BAP), were used at the concentrations, 2, 5, and 10 uM and the negative control was mock infected with 10mM MgCl_2_.

Infection with M11 also caused leaf bending (Figure 4C-D), which was monitored visually and quantified using a leaf bending index, measured as the angle between two lines, one defined by points on the base and mid-point of the petiole and the other by points at mid-point and tip of the leaf (Figure S1B). Significant leaf bending was observed in response to infection with M11 in comparison to mock (Figure 4C), quantified as bending index values of 40.6° and 9.8°, respectively (Figure 4D).. This phenotype was also specific to M11 infection, as the non-host control *T. betulina* had a bending index of 12.1°, which was not significantly different from the mock infected control (Figure 4D).

### Activation of host plant hormone responses

There is evidence of plant hormone production in many *Taphrina* species, including *Taphrina* M11 (see below; Wang et al., 2016). Plant hormones originating from *Taphrina* cells are widely presumed to modulate host plant hormone signalling during pathogenesis; however, this has not been formally tested. To address this, the activation of host hormone transcriptional responses were monitored during infection using two *Arabidopsis* lines transgenically bearing either a cytokinin-responsive (TCR::GUS) or auxin-responsive (DR5::GUS) promoter-reporter construct. These reporter lines underwent various treatments, including infiltration with *Taphrina* strain M11, followed by histological staining of GUS activity to visualize hormone response activation (Figure 5A-B). IAA and BAP were used at three different concentrations as positive controls. The results of the GUS staining experiment showed that both M11 and *T. betulina* were able to activate *Arabidopsis* auxin (Figure 5A) and cytokinin (Figure 5B) transcriptional responses. The IAA response was similar in extent and spatial distribution; localized to the leaf periphery and secondary vasculature in response to both M11 and *T. betulina* (Figure 5A). However, the cytokinin transcriptional response to M11 infection was both stronger and involved more tissues, especially around the base of the leaf, while infection with *T. betulina* resulted in only a small response along the primary leaf vein (Figure 5B).

To determine the role of known plant defence signalling pathways, including immune signalling hormones, a reverse genetics approach was used (Figure 6). Seven knockout mutants were challenged with M11 infection, namely; the jasmonate-insensitive mutants *coi1-16 and jar1*; the *cyp79 b2/b3* double mutant, which is deficient in both indolic glucosinolates and the indole alkaloid phytoalexin, camalexin; the ethylene-insensitive *ein2* mutant; the *pad4* mutant, which is deficient in both camalexin and salicylic acid induction, and the *sid2 and npr1 mutants*, which are deficient in salicylic acid biosynthesis and signalling, respectively. Each mutant was treated both with M11 and mock treatment and evaluated both visually and after trypan blue staining (Figure 6). Upon visual examination, compared to Col-0, *cyp79 b2/b3, coi1-16, pad4,* and *ein2* exhibit enhanced disease susceptibility (EDS) phenotypes expressed as extensive damage and tissue collapse in the leaves inoculated with M11 (Figure 6). The *npr1* and *sid2* mutants had more moderate EDS, with *sid2* showing enhanced leaf deformations. These symptoms did not always correlate with increased cell death, as only *ein2* and *cyp79 b2/b3* exhibited strongly increased cell death that spread out of the infected half of the leaf (Figure 6).

**Figure 6.**
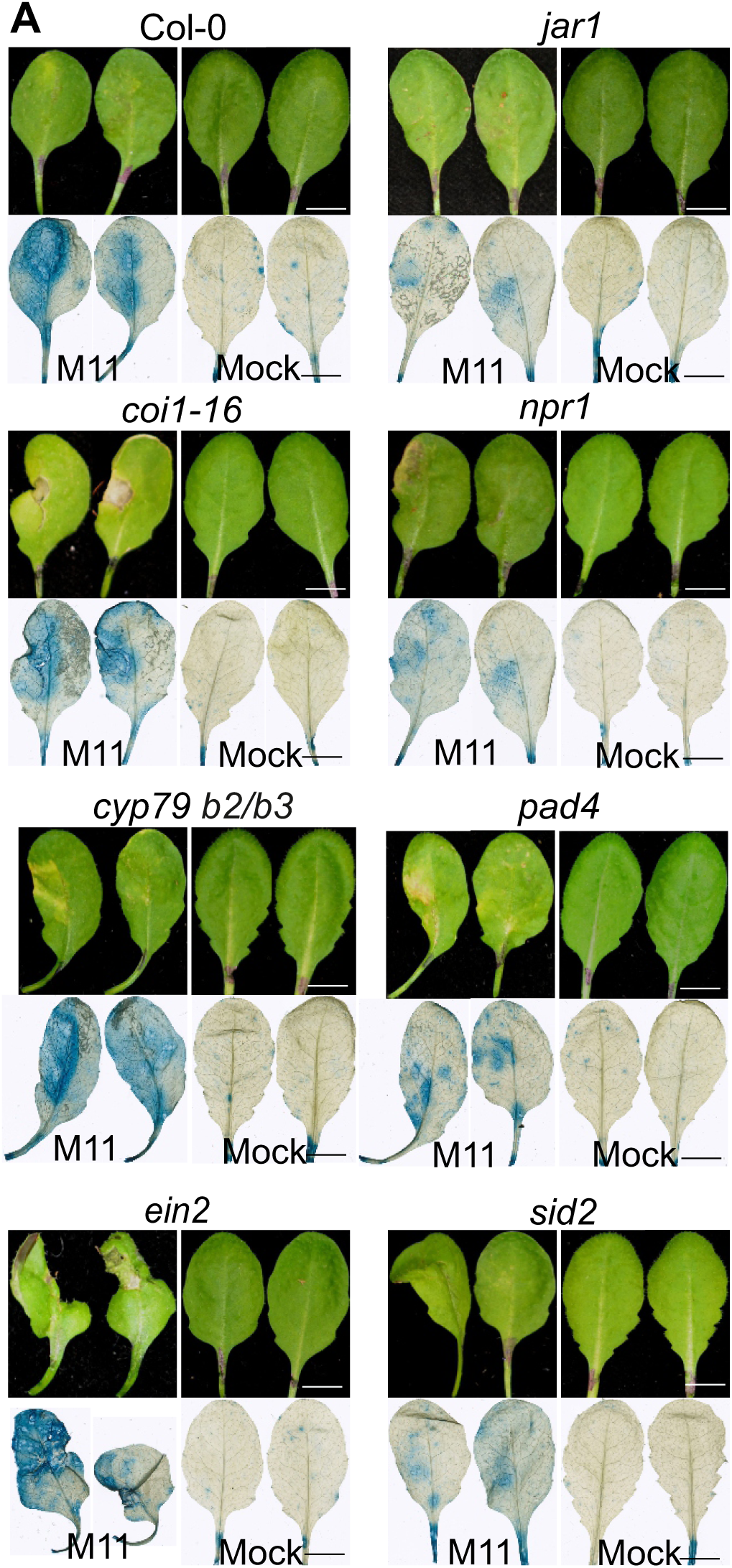
M11 infection of defence signalling mutants. Seven known plant immune signalling mutants were infected with M11 and symptoms were evaluated visually and by visualizing cell death with trypan blue staining. Genotypes used were: wild type Col-0 accession; the jasmonate insensitive *jar1*-1 and *coi1*-16 mutants; *npr1* and *sid2*, which are deficient in salicylic acid perception and biosynthesis, respectively; *cyp 79 b2/b3* double mutant deficient in both indolic glucosinolates and camalexin; the ethylene insensitive *ein2* mutant; and *pad4*, which is deficient in the accumulation of salicylic acid and camalexin. **(A)** The left hand halves of leaves from four-week-old soil-grown plants were hand infiltrated with suspensions of M11 cells or mock infected with 10 mM MgCl_2_ and then photographed and trypan blue stained seven days post infection. All scale bars = 1cm. All leaves were photographed individually against a black background or mounted on microscope slides and the composite figure was assembled in Corel Draw. **(B)** Infections of sterile immune signalling mutants with *Taphrina* M11. *Taphrina* M11 was sprayed onto 24-day-old sterile plants. Yeast growth was quantified through cultivation based technique after nine days. Combined data from three independent experiments is presented; data points from each experiment are represented with a different shape. Pairwise comparison of means (α = 0.05, Tukey test, linear mixed model with biological repeat as a random effect) indicated no differences among genotypes. All scale bars = 1cm. The abbreviation *cyp79\**denotes the *cyp79 b2/b3* double mutant.

### *Taphrina* M11 genome assembly and protein annotations

In order to gain insight into its biology, the genome of *Taphrina* strain M11 was sequenced, resulting in a high quality draft genome assembly of 13.6 Mbp in 234 contigs (Table 1). Characteristics of the *Taphrina* M11 genome were consistent with those of other sequenced *Taphrina* species (Table S3). A total of 5808 proteins were annotated, which is similar to proteomes identified in other previously sequenced *Taphrina* species (Table S1). Ortholog distribution was monitored across M11 and several other species of *Taphrina*, for which genome data is available (Figure 7): additionally several species of the closely related genus *Protomyces* were used for comparison, and *S. pombe* was used as an outgroup to define Taphrinomycotina specific and conserved eukaryotic proteins. On average 38.5% of all proteins were common across the subphylum Taprhinomycotina; however these were not specific to the Taphrinomycotina as they included conserved eukaryotic housekeeping genes. The genera *Taphrina* and *Protomyces* shared 14.9% of their genes, while 5.3% were unique to the genus *Taphrina* (Figure 7). The genus *Protomyces* had slightly more genus specific proteins *-* 7.1%. Only 151 proteins (2.6%) were found to be unique to *Taphrina* strain M11. *Taphrina* M11 shared a sizable portion (132 proteins, 2.3%) of orthologous proteins with *Taphrina* species pathogenic on *Prunus* species, but not with the *T. populina*, which is pathogenic on *Populus* (20 proteins, 0.3%).

**Figure 7.**
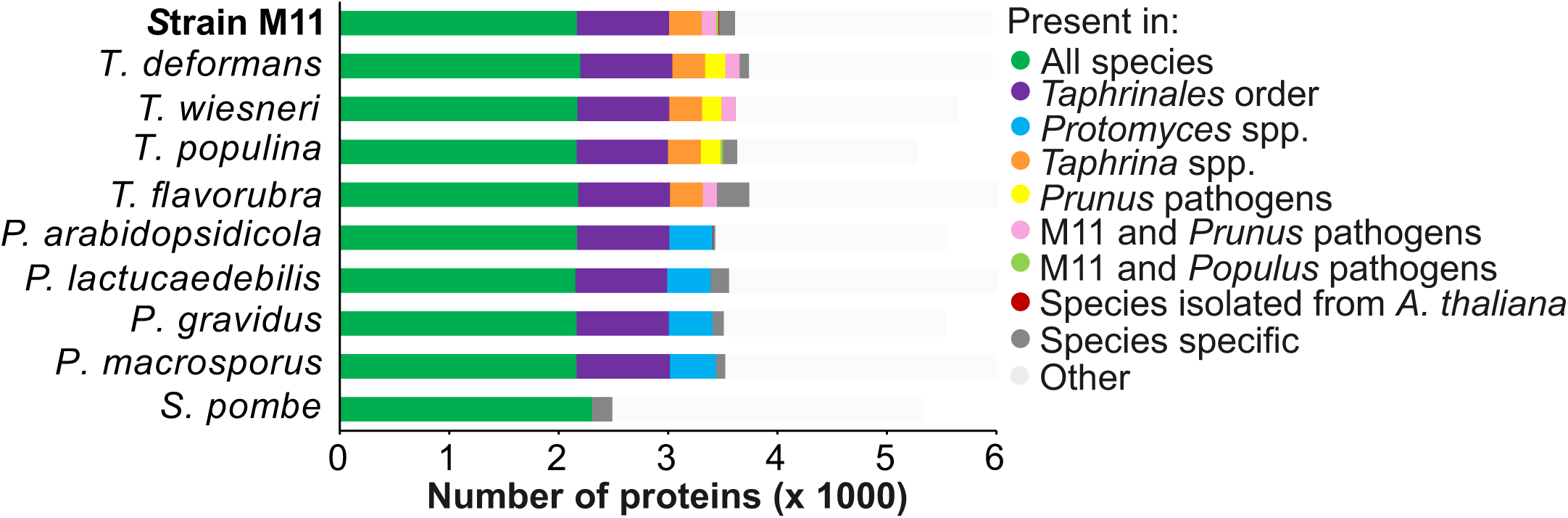
Ortholog distribution in M11 and other Taphrinomycotina yeasts. Ortholog analysis was performed using OrthoVenn2, with E < 0.01 cut-off for ortholog calling. Fission yeast (*Schizosaccharomyces pombe*) was used as an outgroup.

**Table 1.** Taphrina strain M11 genome assembly statistics. Genome assembly quality was analyzed using QUAST tool, version 5.0. For additional explanation of QUAST output see (http://quast.sourceforge.net/).

|  |  |
| --- | --- |
| BioProject<sup>a</sup> | PRJNA487587 |
| BioSample<sup>a</sup> | SAMN09906266 |
| SRA<sup>a</sup> | SRX4936057 |
| Total length (bp) | 13 601 285 |
| Number of contigs (≥ 0 bp) | 382 |
| Total length of contigs (≥ 0 bp) | 13 654 843 |
| Number of contigs (≥ 1000 bp) | 147 |
| Total length of contigs (≥ 1000 bp) | 13 544 983 |
| Total number of contigs | 234 |
| Largest contig (bp) | 514 056 |
| Average GC (%) | 48.83 |
| N50 | 277 880 |
| N75 | 157 923 |
| L50 | 19 |
| L75 | 34 |
| # N's per 100 kbp | 0.02 |
| Annotated genes/orfs | 6496 |
| Number of orfs >100 aa | 14 561 |
| Density (no. of orfs >100 aa/kbp) | 1.07 |

<sup>a</sup>Sequencing data is available at the NCBI under the given accession numbers.

### Candidate effector-like proteins and plant-associated conserved domains

To identify candidate effector-like proteins in M11 genome, a total of 18 660 short (80-333 aa) ORFs were identified, 767 of which contained a secretion signal and were defined as short secreted proteins (SSPs; Table 2A). Of the SSPs, 337 contained 4 or more cysteine residues and were defined as cysteine-rich SSPs (CSSPs). This number of CSSPs was consistent with those present in genomes of other *Taphrina* species (Table 2A). Conserved domains present in these CSSPs were identified (Table 2B). Notably, no LysM domain containing SSPs and CSSPs were detected - this domain is common in effectors from chitin-containing fungi (Table 2B).

**Table 2.**
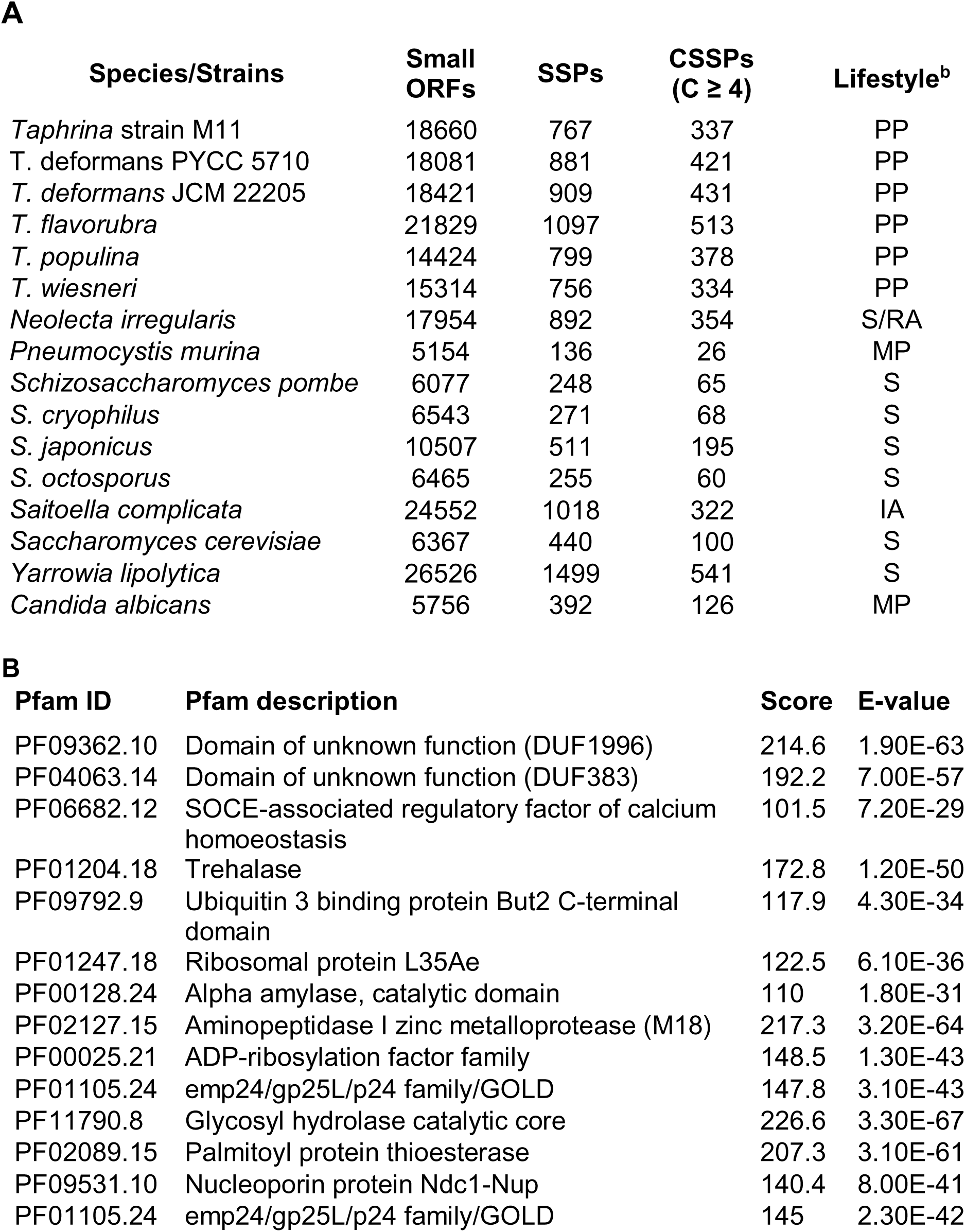
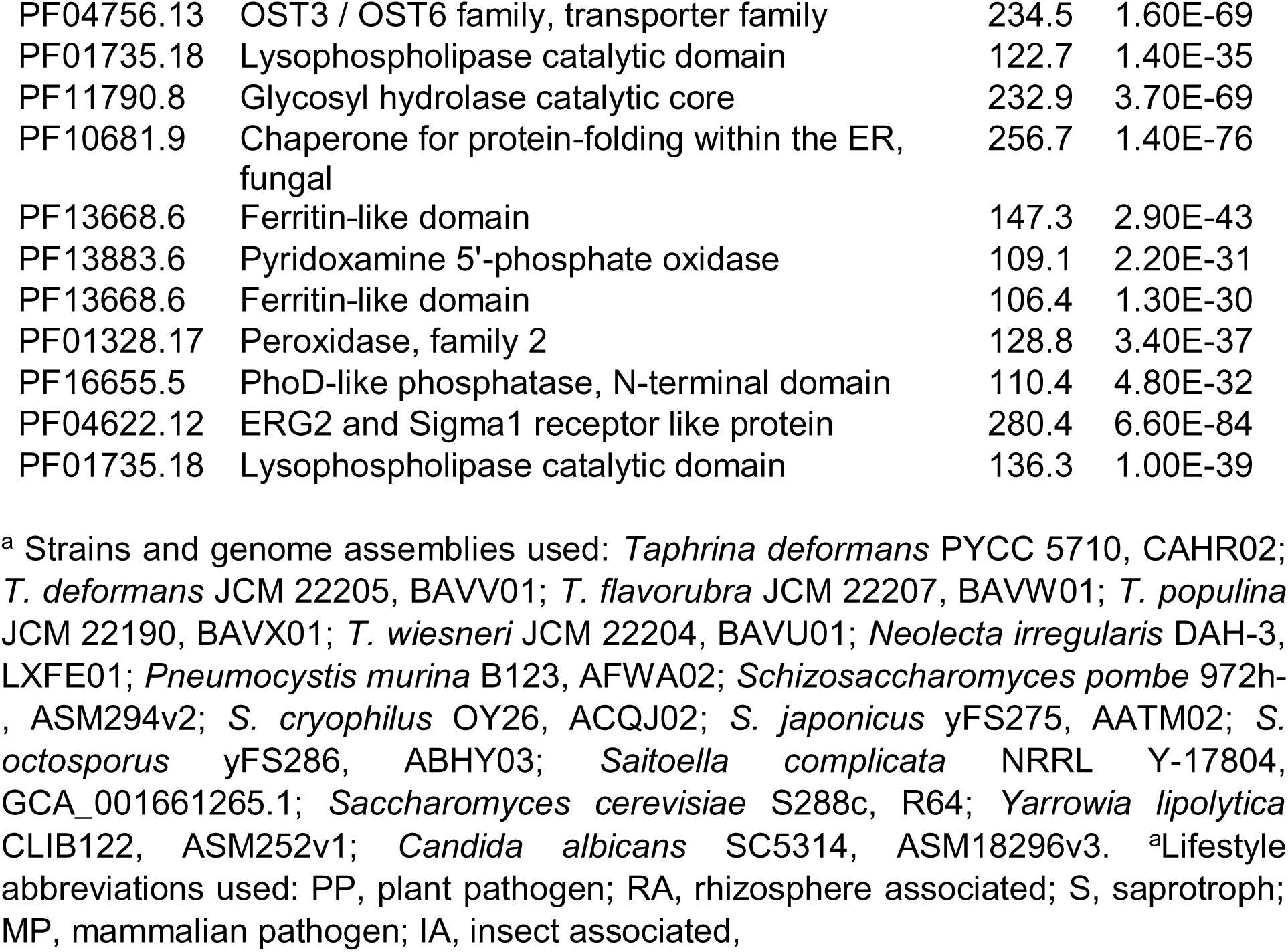
Candidate effector proteins. **(A)** Comparison of identified candidate effector proteins in *Taphrina* strain M11 and other fungi from subphyla Taphrinomycotina and Saccharomycotina^a^. Number of open reading frames (ORFs), short secreted proteins (SSPs) and cysteine rich SSPs (CSSPs) is shown. **(B)**The conserved protein domains were identified from *Taphrina strain* M11 SSPs using HMMER v3.2.1 (E value 1e-30) homology searches against the Pfam database. The presented E-values and scores are for full sequences.

**A**
| Species/Strains | Small ORFs | SSPs | CSSPs (C ≥ 4) | Lifestyle<sup>b</sup> |
| --- | --- | --- | --- | --- |
| *Taphrina* strain M11 | 18660 | 767 | 337 | PP |
| *T. deformans* PYCC 5710 | 18081 | 881 | 421 | PP |
| *T. deformans* JCM 22205 | 18421 | 909 | 431 | PP |
| *T. flavorubra* | 21829 | 1097 | 513 | PP |
| *T. populina* | 14424 | 799 | 378 | PP |
| *T. wiesneri* | 15314 | 756 | 334 | PP |
| *Neolecta irregularis* | 17954 | 892 | 354 | S/RA |
| *Pneumocystis murina* | 5154 | 136 | 26 | MP |
| *Schizosaccharomyces pombe* | 6077 | 248 | 65 | S |
| *S. cryophilus* | 6543 | 271 | 68 | S |
| *S. japonicus* | 10507 | 511 | 195 | S |
| *S. octosporus* | 6465 | 255 | 60 | S |
| *Saitoella complicata* | 24552 | 1018 | 322 | IA |
| *Saccharomyces cerevisiae* | 6367 | 440 | 100 | S |
| *Yarrowia lipolytica* | 26526 | 1499 | 541 | S |
| *Candida albicans* | 5756 | 392 | 126 | MP |

**B**
| Pfam ID | Pfam description | Score | E-value |
| --- | --- | --- | --- |
| PF09362.10 | Domain of unknown function (DUF1996) | 214.6 | 1.90E-63 |
| PF04063.14 | Domain of unknown function (DUF383) | 192.2 | 7.00E-57 |
| PF06682.12 | SOCE-associated regulatory factor of calcium homoeostasis | 101.5 | 7.20E-29 |
| PF01204.18 | Trehalase | 172.8 | 1.20E-50 |
| PF09792.9 | Ubiquitin 3 binding protein But2 C-terminal domain | 117.9 | 4.30E-34 |
| PF01247.18 | Ribosomal protein L35Ae | 122.5 | 6.10E-36 |
| PF00128.24 | Alpha amylase, catalytic domain | 110 | 1.80E-31 |
| PF02127.15 | Aminopeptidase I zinc metalloprotease (M18) | 217.3 | 3.20E-64 |
| PF00025.21 | ADP-ribosylation factor family | 148.5 | 1.30E-43 |
| PF01105.24 | emp24/gp25L/p24 family/GOLD | 147.8 | 3.10E-43 |
| PF11790.8 | Glycosyl hydrolase catalytic core | 226.6 | 3.30E-67 |
| PF02089.15 | Palmitoyl protein thioesterase | 207.3 | 3.10E-61 |
| PF09531.10 | Nucleoporin protein Ndc1-Nup | 140.4 | 8.00E-41 |
| PF01105.24 | emp24/gp25L/p24 family/GOLD | 145 | 2.30E-42 |
| PF04756.13 | OST3 / OST6 family, transporter family | 234.5 | 1.60E-69 |
| PF01735.18 | Lysophospholipase catalytic domain | 122.7 | 1.40E-35 |
| PF11790.8 | Glycosyl hydrolase catalytic core | 232.9 | 3.70E-69 |
| PF10681.9 | Chaperone for protein-folding within the ER, fungal | 256.7 | 1.40E-76 |
| PF13668.6 | Ferritin-like domain | 147.3 | 2.90E-43 |
| PF13883.6 | Pyridoxamine 5'-phosphate oxidase | 109.1 | 2.20E-31 |
| PF13668.6 | Ferritin-like domain | 106.4 | 1.30E-30 |
| PF01328.17 | Peroxidase, family 2 | 128.8 | 3.40E-37 |
| PF16655.5 | PhoD-like phosphatase, N-terminal domain | 110.4 | 4.80E-32 |
| PF04622.12 | ERG2 and Sigma1 receptor like protein | 280.4 | 6.60E-84 |
| PF01735.18 | Lysophospholipase catalytic domain | 136.3 | 1.00E-39 |

<sup>a</sup> Strains and genome assemblies used: *Taphrina deformans* PYCC 5710, CAHR02; *T. deformans* JCM 22205, BAVV01; *T. flavorubra* JCM 22207, BAVW01; *T. populina* JCM 22190, BAVX01; *T. wiesneri* JCM 22204, BAVU01; *Neolecta irregularis* DAH-3, LXFE01; *Pneumocystis murina* B123, AFWA02; *Schizosaccharomyces pombe* 972h-, ASM294v2; *S. cryophilus* OY26, ACQJ02; *S. japonicus* yFS275, AATM02; *S. octosporus* yFS286, ABHY03; *Saitoella complicata* NRRL Y-17804, GCA\_001661265.1; *Saccharomyces cerevisiae* S288c, R64; *Yarrowia lipolytica* CLIB122, ASM252v1; *Candida albicans* SC5314, ASM18296v3. <sup>a</sup>Lifestyle abbreviations used: PP, plant pathogen; RA, rhizosphere associated; S, saprotroph; MP, mammalian pathogen; IA, insect associated,

We also queried all the *Taphrina* M11 annotated proteins for conserved domains previously identified as being typical of plant-associated microorganisms by (Levy et al., 2017). Proteins containing RNA recognition motif were common in the M11 genome (Table S5), which is a typical characteristic of biotrophic pathogens (Pandaranayaka et al., 2019). The most common domain, however, was the protein tyrosine kinase domain.

### Putative plant hormone biosynthesis pathways

Genes involved in indole acetic acid (IAA) and cytokinin production (Table 4) were identified in the M11 genome using known biosynthesis genes from other species, which represented four possibly routes for IAA biosynthesis in microbes; specifically, the indole-3-acetamide (IAM), indole-3-pyruvate (IPyA), indole-3-acetonitrile (IAN), and tryptamine (TAM) pathways. Remarkably, the *Taphrina* M11 genome contained complete enzymatic machinery for IAA production via three different routes – the IAM, IPyA, and TAM pathways (Table 4). Enzymes involved in these pathways were also conserved in *T. deformans* (Table S6).

Two key enzymes of cytokinin biosynthesis were also identified in M11 genome (Table 4) - tRNA-isopentenyltransferase and cytokinin phosphoribohydrolase. However, the presence of other enzymes involved in this pathway cannot be excluded, as no query sequences for other steps of the pathway were available from closely related fungi.

### Biosynthesis of potentially immunoactive cell wall polysaccharides

Microbial cell walls are a major source of PAMPs such as chitin and linear β-1,3-glucan, which are recognized by plant immune receptors. To predict potential PAMPs present in *Taprhina* M11 cell walls we looked for putative cell wall biosynthesis proteins in *Taphrina* M11 genome and compared them with biochemical evidence for presence of different cell wall polysaccharides from other studies (Table 3). Despite biochemical studies indicating that yeast cells of *Taphrina deformans* contain little to no chitin (Petit and Schneider, 1983), we identified two conserved chitin synthases in the *Taphrina* M11 genome and three conserved chitin synthases in *T. deformans* (Figures S3, S4, and S5). Additionally, a putative chitin deacetylase homolog was found, which could potentially be used by M11 to transform chitin into chitosan (Table 3).

**Table 3.**
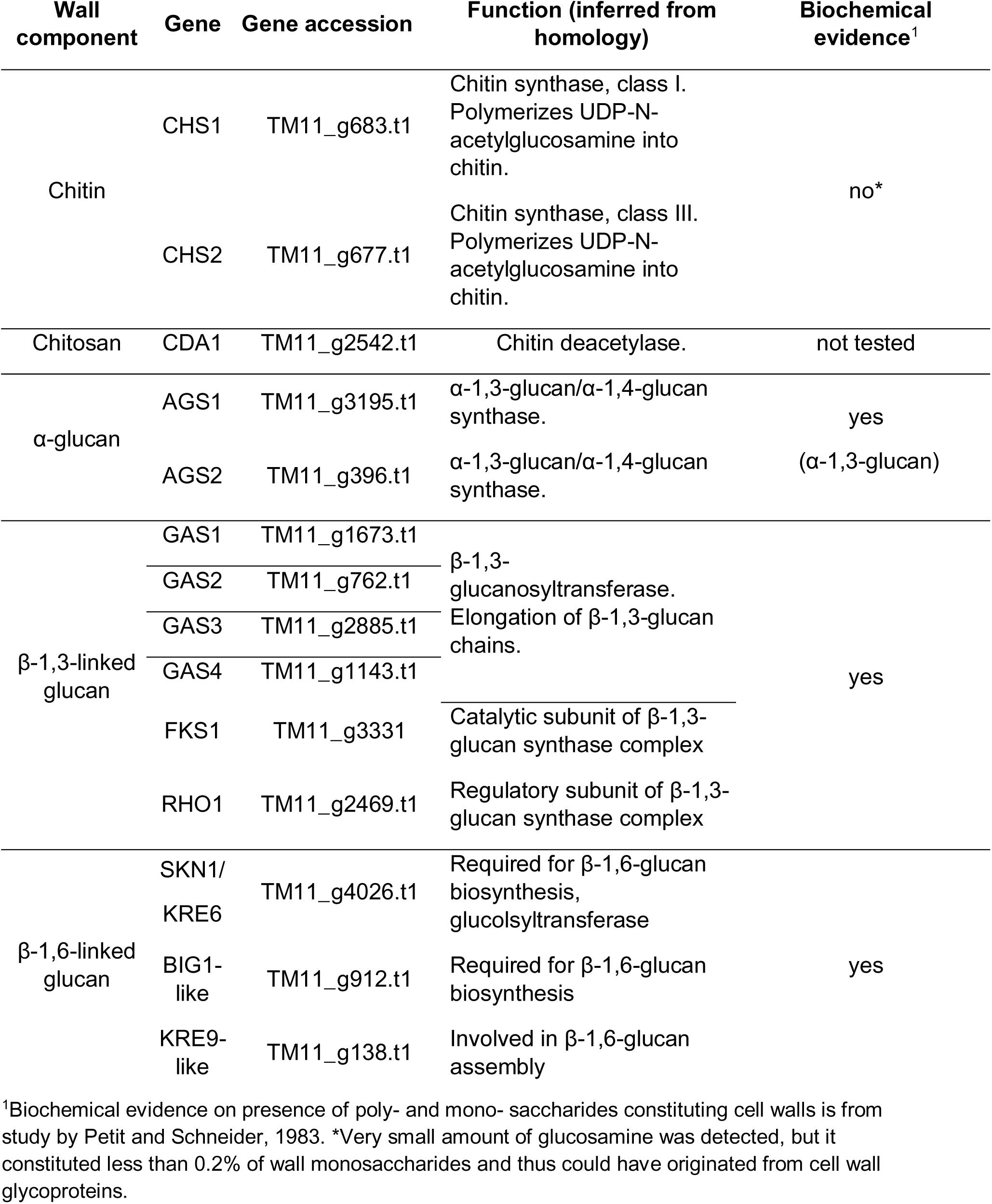
Putative cell wall biosynthesis genes in Taphrina strain M11. Potential cell wall components were predicted based on the putative cell wall biosynthesis genes and biochemical evidence presented in Petit and Schneider, 1983. Sequences of well-described homologs from *S. cerevisiae, S. pombe,* and *A. nidulans* were used as protein blast queries. Additionally, all predicted genes containing conserved protein domains specific for the cell wall biosynthesis genes were analyzed to confirm their identity. For additional information on query sequences, putative homolog sequences, and protein blast results see Tables S6 and S7.

**Table 4.**
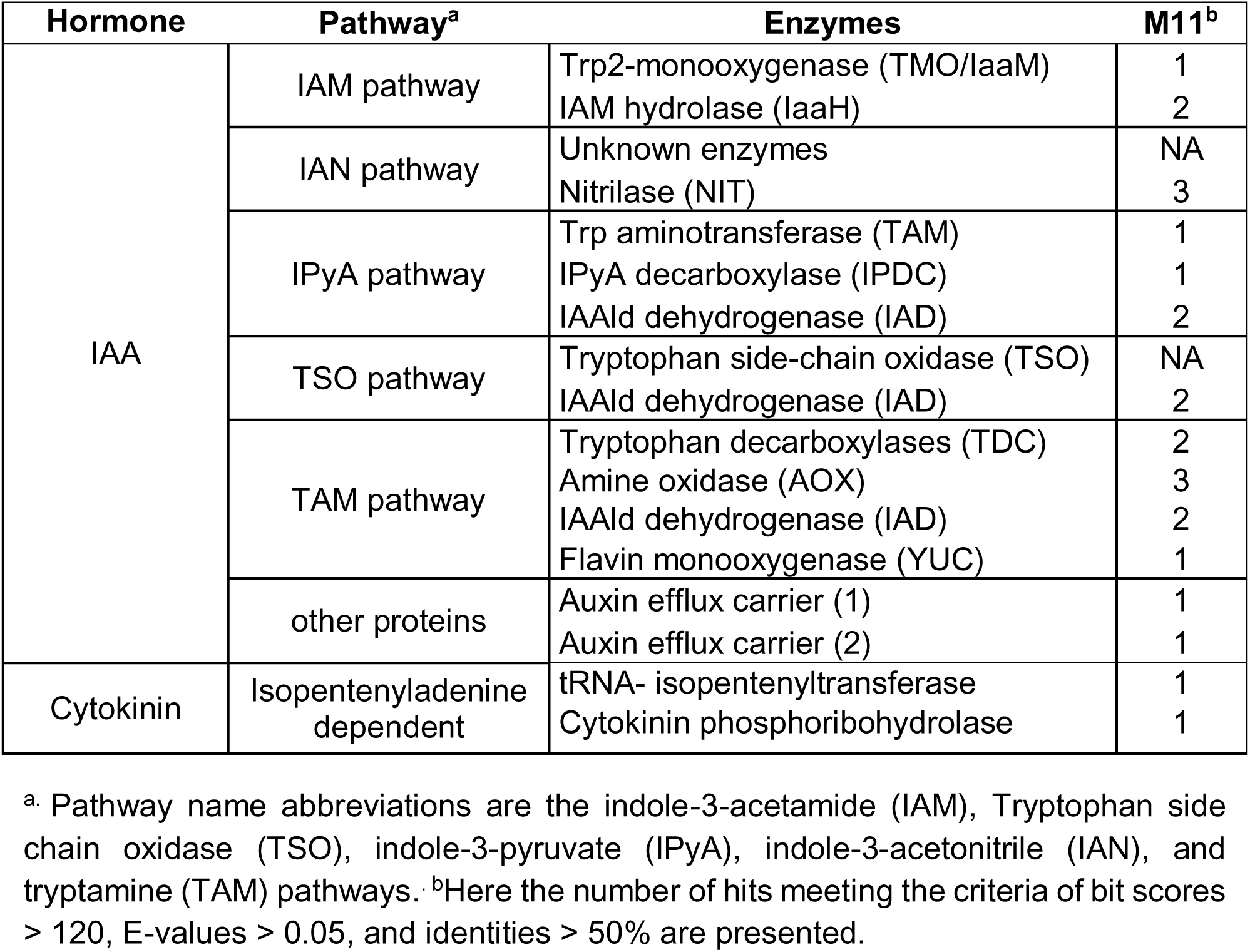
Auxin and cytokinin biosynthesis pathways in *Taphrina* strain M11.

The M11 genome contains homologs of proteins necessary for production of β-glucans both with β-1,3-linkages and β-1,6-linkages (Table 3). The presence of these polysaccharides was also supported by previous biochemical analysis (Petit and Schneider, 1983). Another glucose polysaccharide potentially present in M11 cell walls is α-glucan. We identified two α-1,3-glucan/α-1,4-glucan synthase genes in M11 genome (Table 3). According to biochemical evidence, similar to *Schizosaccharomyces pombe Taprhina* species contain predominantly α-1,3-glucan (Petit and Schneider, 1983).

## Discussion

### M11 causes disease on Arabidopsis

Infection of *Arabidopsis* with the species *T. betulina*, a *Taphrina* not adapted to *Arabidopsis* -- used here as a control for the non-host defence response, resulted in accumulation of H_2_O_2_, activation of rapid HR-like cell death, no growth *in planta,* and no late leaf deformation or other symptoms. These results indicate that *Arabidopsis* has immunity against *T. betulina* and likely other *Taphrina* species. However, the activation of HR-like cell death observed here was in contrast to most forms of non-host resistance (Lee et al., 2017). This type of non-host resistance appears to share similarities to effector triggered immunity (ETI), normally conditioning host immunity to adapted pathogens. There are several known examples of this; however, the mechanisms and roles of ETI-like resistance in non-host resistance remain poorly understood (Stam et al., 2014); (Lee et al., 2017). Infections with *Taphrina* strain M11, which was isolated from wild *Arabidopsis* in the field, resulted in an attenuated ROS response compared to *T. betulina*, a small amount of cell death that was consistent with symptom development, successful multiplication on *Arabidopsis*, and leaf symptom development including chlorosis and late presentation of leaf deformations. These findings support that strain M11 is adapted to, and pathogenic on *Arabidopsis*. M11 having the lifestyle of a plant pathogen was also supported by several of its genomic features, such as; a full complement of candidate effector proteins, similar to those found in other *Taphrina* species; and the presence of conserved protein domains previously shown to be specific to plant associated microbes. There are currently no resistance mechanisms against *Taphrina* known at the mechanistic level; although several studies have begun to address the possibility of such resistance. Evidence of genetic resistance against witches’ broom disease caused by *T. betulina* segregating in natural populations of birch (Christita and Overmyer, 2021). *Taphrina* resistance has also been addressed in peach (Goldy et al., 2017), where *Taphrina* causes significant economic losses and chemical fungicides are the only available means of control.

*Taphrina* strain M11 was found to have an ITS sequence that was 99.% similar to the ITS of *T. tormentillae* (Wang et al., 2016); exceeding the 98.4% threshold determined for delimitation of yeast species (Vu et al., 2016). This suggests that M11 is a strain of *T. tormentillae*. However, as has been previously noted, ITS sequences do not always offer good resolution to the species level in the genera *Taphrina* and *Protomyces* (Rodrigues and Fonseca, 2003; Wang et al., 2021). Further studies with other phylogenetic markers will be required before the relationship between M11 and *T. tormentillae* can be determined. The *actin1* gene was found to function well as a lineage specific secondary phylogenetic marker for species determination in the closely related genus *Protomyces* (Wang et al., 2021). Although it remains to be tested, this nuclear gene DNA marker may also function well for the genus *Taphrina*. Remarkably, the two other tested *Taphrina* strains --*T. tormentillae* strain PYCC 5727 and the closely related strain CBS 332.55, which is also most likely a strain of *T. tormentillae*--were also able to grow on *Arabidopsis,* but to slightly lower levels. Further infections with these two strains will be required to determine if pathogenicity on *Arabidopsis* is a common feature of all strains related to *T. tormentillae.* Reciprocal infections of these three strains on *Potentilla* species known to be hosts for *T. tormentillae* will also be required in future studies to fully address host range of this *Taphrina* species.

### Plant defenses, candidate effectors, and yeast PAMPs

Infection of known immune signaling mutants of *Arabidopsis* suggested that ethylene and jasmonate signaling are required for resistance against *Taphrina* M11. Previous studies with other related systems have shown similar results. A strain (strain C29) of *Protomyces*, the sister genus to *Taphrina*, was isolated from *Arabidopsis* and named *P. arabidopsidicola* (Wang et al., 2021). Strain C29 was not pathogenic, but persisted in the *Arabidopsis* phyllosphere and activated enhanced immunity against the fungal necrotrophic pathogen *Botrytis*, which was associated with activation of MAPK signalling and markers of camalexin biosynthesis and salicylic acid signalling (Wang et al., 2019) Additionally, treatment of plants with yeast cell wall preparations, yeast extract, or autoclave killed *S. cerevisiae* have been shown to activate resistance and activate defence signalling pathways (Raache et al., 2006; Khokon et al., 2010). The above results indicate that yeast induce defence signalling pathways similar to those responding to other pathogenic fungi. However, yeasts and dimorphic yeast-like fungi present a different set of surface molecules compared to other fungi and the major yeast PAMPs remain uncharacterized.

To address the possibility of unique PAMPs in *Taphrina*, analysis of potential cell wall carbohydrate biosynthesis genes in the M11 genome was undertaken. Although conserved chitin synthases were found in M11 and *T. deformans* genome (Figures S3-S5), biochemical studies done in the past on *Taphrina* and the closely related *Protomyces* cell walls do not indicate the presence of chitin (Petit and Schneider, 1983; Valadon et al., 1962). However, in those studies the cell walls were isolated only from the yeast-like cells of *Taphrina* and *Protomyces*. As indicated by studies on other dimorphic fungi, cell wall composition can change depending on the growth form and hyphae can contain substantially more or less chitin than the yeast-like cells (Díaz-Jiménez et al., 2012). Also in the cell walls of *S. pombe* - the best studied yeast from the subphylum *Taphrinomycotina* - chitin can only be detected in ascospores (Arellano et al., 2000). Thus, the presence of conserved chitin synthases in *Taphrina* M11 and *T. deformans* genomes suggests that *Taphrina* could have detectable chitin in other growth forms or produce it only during ascospore formation like *S. pombe*.

Notably, similar to the closely related species from *Protomyces* genus (Wang et al., 2019), *Taphrina* M11 genome lacks effector-like proteins containing LysM domain. LysM domain containing effectors are a common strategy among fungi for hiding chitin from plant pattern recognition receptors (Gong et al., 2020). The lack of LysM effectors suggests that chitin is of lesser importance in host interactions or that *Taphrina* yeasts might use different strategies for masking chitin such as non-LysM effectors, chitin deacetylation, or chitin inaccessibility mediated by layers of other polysaccharides, such as β-glucans or non-degradable α-1,3-glucan. In support of the latter two potential strategies, a putative chitin deacetylase and two α-glucan synthases were found in M11 genome.

β-glucans are known to elicit immune responses in a wide range of plants (Fesel and Zuccaro, 2016), including *Arabidopsis* (Melida et al., 2018; Rebaque et al., 2021). In *Taphrina* strain M11 we identified the necessary machinery for production of β-glucans with β-1,3- and β-1,6-linkages. The presence of these polysaccharides in *Taphrina* is also supported by biochemical studies (Petit and Schneider, 1983). Interestingly, *Taphrina* M11 genome contains only one β-1,3-glucan synthase FKS homolog. *S. pombe,* which has a similar cell wall composition to *Taphrina* (Perez et al., 2018), has four FKS homologs with different functions and regulation. In the dimorphic plant pathogen *Ustilago maydis* and other species having only one FKS homolog, the gene is essential for survival and is constitutively expressed (Ruiz-Herrera and Ortiz-Castellanos, 2019).

Apart from chitin and β-1,3-glucans that have been described to trigger immune response in *Arabidopsis* (Xue et al., 2019; Melida et al., 2018), cell walls of M11 could contain glycoproteins with immunoactive glycans. From cell wall monosaccharide analysis, it is known that *Taphrina* walls contain mannose, rhamnose, and galactose (Petit and Schneider, 1983; Sjamsuridzal et al., 1997). These could be arranged in galactomannans and rhamnomannans as seen in the filamentous plant-pathogenic ascomycete *Rhynchosporium secalis* (Pettolino et al., 2009). Both galactomannans and rhamnomannans are recognized by innate immunity of animals (Barreto-Bergter and Figueiredo, 2014) and could possibly be also recognized by plants. *Taphrina* cell walls could also contain simple mannose polysaccharides. Mannopeptides (protease digested mannan glycoproteins) have been previously demonstrated to elicit immune response in tomato cell cultures (Grosskopf et al., 1991).

### Plant Hormones in plant-Taphrina interactions

*Taphrina* are well documented as producers of the plant hormones auxin and cytokinin (Kern and Naef-Roth, 1975; Cissé et al., 2013; Tsai et al., 2014; Streletskii et al., 2016, Streletskii et al., 2019; Wang et al., 2016). Auxin and cytokinin production is widely believed to be responsible for the dramatic leaf deformation and tumour symptoms typical of the plant pathogens belonging to the genus *Taphrina*, although this has never been formally tested. Previous studies have examined *in vitro* plant hormone production by *Taphrina* species, including *Taphrina* Strain M11. Here, taking advantage of the genetic tools available with the model plant *Arabidopsis*, activation of plant hormone response was monitored during infection with M11*Taphrina*. Auxin response was activated slightly and non-specifically, in response to both pathogenic and non-pathogenic *Taphrina* species. In *Ustilago maydis*, a pathogen that shares many similarities with the *Taphrina*, reduced auxin production by 60% by loss of function in multiple biosynthesis genes had no effect on tumour formation during infection of its host maize, suggesting that auxin may not be required for tumour formation. In contrast to IAA responses, *Arabidopsis* cytokinin response was specifically activated in response to the pathogenic M11 strain, but not the non-host control. Cytokinins are key plant developmental hormones that promote cell divisions (Argueso et al., 2009) and are also known to be produced by several plant associated microorganisms, including other tumor inducing pathogens (Pertry et al., 2010). In Ustilago, cytokinin production was required for full induction of tumors (Morrison et al., 2015). Several studies have presented evidence of cytokinin production in *Taphrina* species (Sommer, 1961; Barthe and Bulard, 1974; Streletskii et al., 2019). Further studies will be required to test the role of auxin and assess if cytokinins from M11 *Taphrina* are associated with phenotypic changes in response to pathogenic infections.

To further explore the ability for plant hormone production in *Taphrina*, the auxin and cytokinin biosynthesis genes were examined the M11 genome and revaluated from the genome of *T. deformans*. Both of these*Taphrina* species have the necessary enzymes for auxin production through three different pathways - IAM pathway, IPyA pathway, and TAM pathway (Table 4, Table S6). To our knowledge, this is the first study in which tryptophan monooxygenase and indole acetamide hydrolase (IAM pathway) have been found in *Taphrina* genomes. In previous studies enzymes involved in IAM pathway have not been detected in *Taphrina* species (Tsai et al, 2014). In another study, only *TAM* and *LAD* genes of the IPyA pathway were found in the genome of *T. deformans* (Cissé et al., 2013). In comparison, the closely related *Protomyces* species, which have been demonstrated to secrete auxin into culture media, have only one conserved IAA biosynthesis pathway - IPyA pathway (Wang et al., 2019). These results raise the question of the need for these multiple IAA biosynthesis pathways and their function. A model for future testing is suggested by the multiple roles played by auxin in microbes (Spaepen and Vanderleyden, 2011). Auxin is used by pathogens to subvert host immunity and promote infection (Naseem and Dandekar, 2012; Ma and Ma, 2016; Fu and Wang, 2011); in beneficial microbes to promote host growth (Ahmad et al., 2008; Contreras-Cornejo et al., 2009). Additionally, auxin functions in fungal development (Chanclud and Morel, 2016), and is an important adaptation promoting survival in the phyllosphere (Vorholt, 2012; Kemler M., 2017). The various IAA biosynthesis pathways in *Taphrina* species may be functionally divergent as has been seen in bacterial systems. *Pantoea agglomerans* pv gypsophilae has dual IAA biosynthesis pathways that are differentially expressed and differentially required; the IAM pathway for pathogenesis, infection and gall formation and the IPyA pathway for fitness in the phyllosphere (Manulis et al., 1998; Chalupowicz et al., 2009).

In *Taphrina* M11 we identified key enzymes necessary for cytokinin production, thus further supporting the hypothesis that observed increase in cytokinin levels of arabidopsis might be coming from yeast-produced cytokinin (Table 4).

## Acknowledgement

Authors thanks to Tuomas Puukko, and Leena Grönholm, for excellent technical support. MC is a member of the University of Helsinki Doctoral Program in Plant Science (DPPS); AA is a member of the University of Helsinki Microbiology and Biotechnology Doctoral Program (MBDP). This work was supported by the following grants: The Academy of Finland Center of Excellence in Primary Producers 2014-2019 (decisions #271832 and 307335); a research grant to MC from the Finnish Society for Forestry Sciences; a research grant to AA from the Kuopio Nature Friends’ Association (Kuopion Luonnon Ystäväin Yhdistys); research allocations to AA and MC from the University of Helsinki YEB graduate school, the MBDP, and DPPS; a PhD Fellowship from MPDP granted to AA, and a PhD Fellowship from The Indonesian Fund for Education (LPDP) granted to MC. Computing resources provided by the Finnish IT Center for Science (CSC; www.csc.fi) are gratefully acknowledged. The authors declare no conflict of interest related to this work.

## Tables

**Table S1. Ortholog distribution in selected Taphrinomycotina species.** Protein annotations of selected species were analyzed using OrthoVenn2 web platform. *Taphrina* strain M11 and *Protomyces arabidopsidicola* were isolated from wild *Arabidopsis thaliana* plants. Other included *Protomyces* species are known to infect Compositae and Umbelliferae plants. *T. deformans*, *T. wiesneri*, and *T. flavorubra* are pathogenic to *Prunus* species. Host of *T. populina* is *Populus nigra*. *Schizosaccharomyces pombe* is phylogenetically most distant to other species and has a non-pathogenic lifestyle.

**Table S2. Conserved domains in Taphrina strain M11 annotated proteins.** Data was generated using HMMER v3.3 (E value 1e-30) through homology searches against the Pfam database. For the full explanation of HMMER output see the HMMER User’s Guide at www.hmmer.org

**Table S3. Comparison of genome statistics and candidate effector proteins.** Analysis was performed with *Taphrina* strain M11 and other fungi from subphyla Taphrinomycotina and Saccharomycotina. Used strains and genome assemblies: *Taphrina deformans* PYCC 5710, CAHR02; *T. deformans* JCM 22205, BAVV01; *T. flavorubra* JCM 22207, BAVW01; *T. populina* JCM 22190, BAVX01; *T. wiesneri* JCM 22204, BAVU01; *Neolecta irregularis* DAH-3, LXFE01; *Pneumocystis murina* B123, AFWA02; *Schizosaccharomyces pombe* 972h-, ASM294v2; *S. cryophilus* OY26, ACQJ02; *S. japonicus* yFS275, AATM02; *S. octosporus* yFS286, ABHY03; *Saitoella complicata* NRRL Y-17804; *Saccharomyces cerevisiae* S288c, R64; *Yarrowia lipolytica* CLIB122, ASM252v1; *Candida albicans* SC5314, ASM18296v3. Secretion signal was identified using SignalP 4.1 tool.

**Table S4. Conserved domains in short secreted proteins (SSPs).** Table depicts the full data from the searches for the conserved protein domain hits found in SSPs from *Taphrina* strain M11 and reported in Table 2b. Data was generated using HMMER v3.2.1 (E value 1e-30) through homology searches against the Pfam database. For the full explanation of HMMER output see The HMMER User’s Guide at www.hmmer.org

**Table S5. Conserved plant-associated domains in the annotated proteins of *Taphrina* species strain M11.** Plant associated domains were previously defined from plant associated prokaryotes (Levy et al., 2017).

**Table S6. Indole-3-acetic acid (IAA) biosynthesis pathways in two Taphrina species.** Analysis compares IAA biosynthesis genes of *Taphrina* species strain M11 and *T. deformans*. Genes were first identified in strain M11 (Table 4) using potential IAA biosynthesis genes from other fungal species as BLAST search terms, as in (Wang et al., 2019). The genes of IAA biosynthesis enzymes found in M11 genome were then used as search terms with the BLAST tool against the *T. deformans* genome. Pathway name abbreviations are the indole-3-acetamide (IAM), indole-3-pyruvate (IPyA), indole-3-acetonitrile (IAN), and tryptamine (TAM) pathways.

**Table S7. Putative cell wall biosynthesis genes from *Taphrina* M11.** Blast protein search was performed with default search parameters against the *Taphrina* M11 genome using the well characterized cell wall biosynthesis genes from model fungal species as queries. Known conserved protein domains found in such genes were also used as queries. Query genes and conserved domains used are listed in Table S8.

**Table S8: Cell wall biosynthesis query sequences and conserved domains used.** Queries used to identify putative proteins involved in M11 cell wall polysaccharide biosynthesis. Conserved domains (Pfam database) are indicated in blue. Gene accession numbers are from UniProt database. NA - not applicable

---

**Figure S1.**
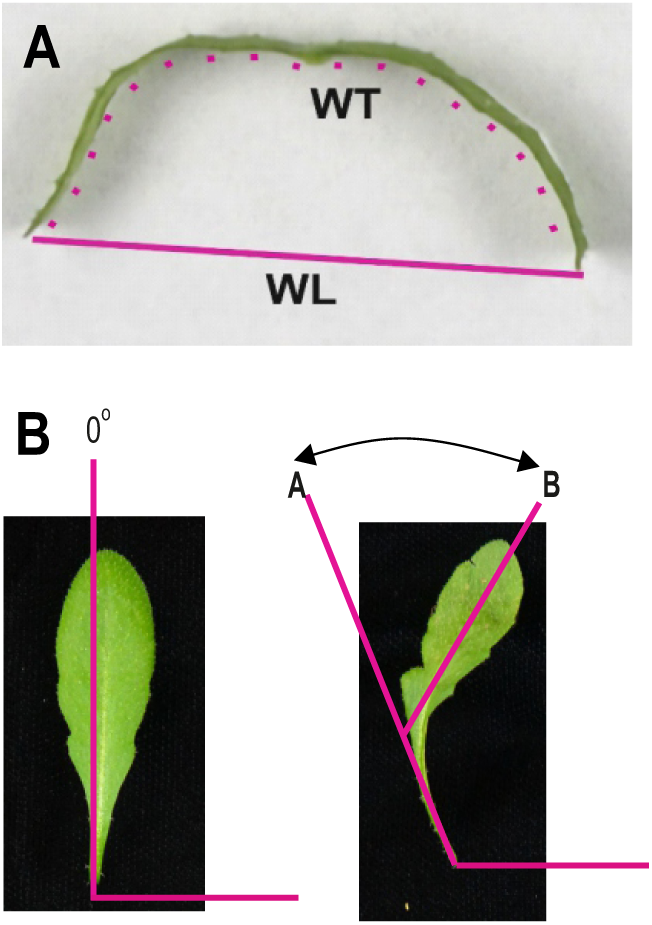
Leaf curling index and leaf bending index measurement. Leaf curling **(A)** was measured as in (Booker et al., 2004), briefly this calculates the ratio of the leaf width (W_L_) to the total width (W_T_), so that leaves with greater curling will have smaller values for the leaf curling index. Leaf bending **(B)** was quantified by measuring the angle between a line following the base of the petiole and a second line defined by points at the middle and tip of the leaf, such that leaves with greater bending will have higher values.

**Figure S2.**
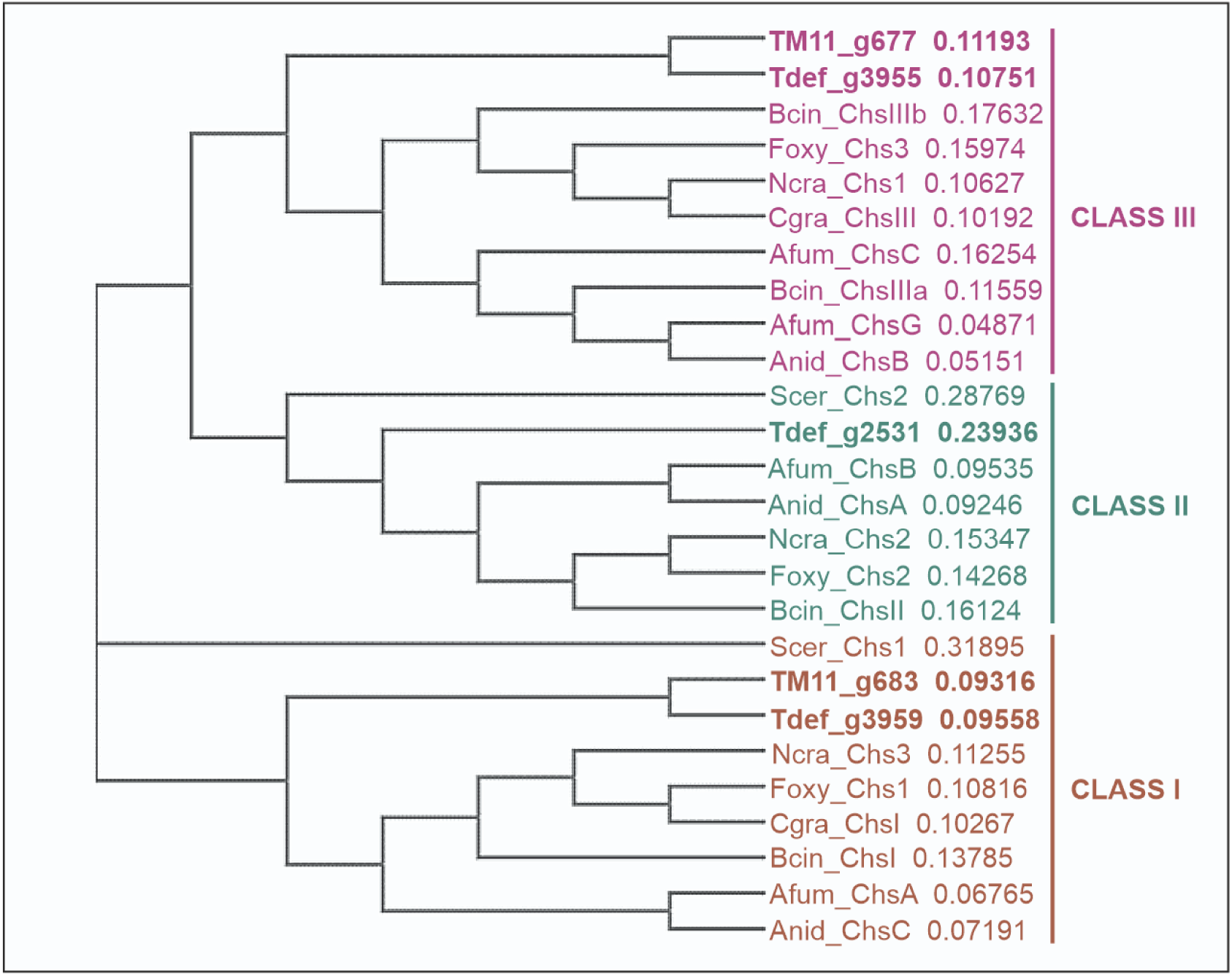
Phylogeny of chitin synthase genes. Chitin synthase genes were identified from *Taphrina* species and fungal model organisms from the phylum Ascomycota and aligned using the Clustal Omega multiple sequence alignment tool. Species abbreviations used: TM11, *Taphrina* species strain M11; Tdef, *Taphrina deformans*, Bcin, *Botrytis cinerea*; Foxy, *Fusarium oxysporum*; Ncra, *Neurospora crassa*, Cgra, *Colletotrichum graminicola*; Afum, *Aspegillus fumigatus*; Anid, *Aspegillus nidulans*.

**Figure S3.**
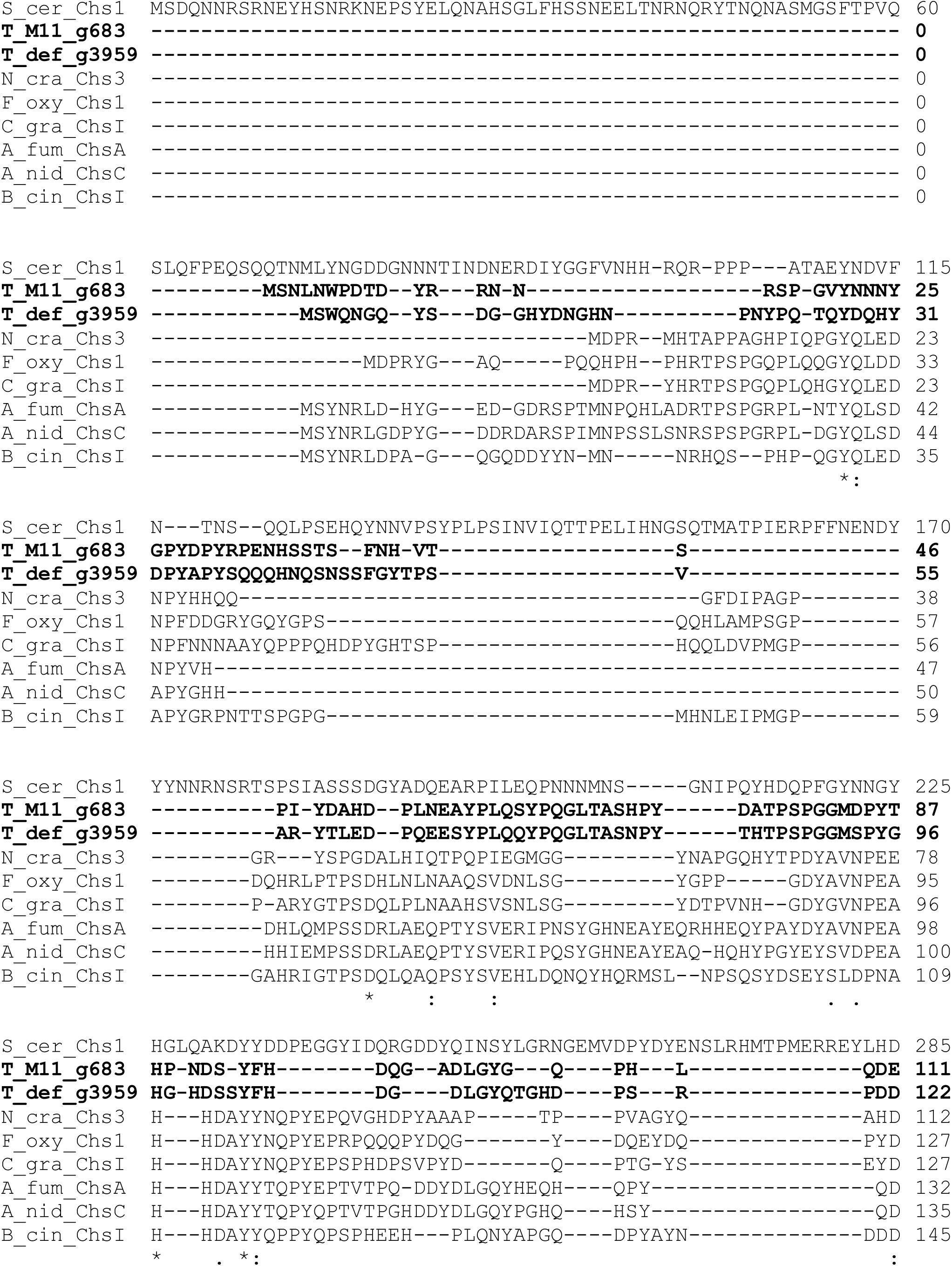
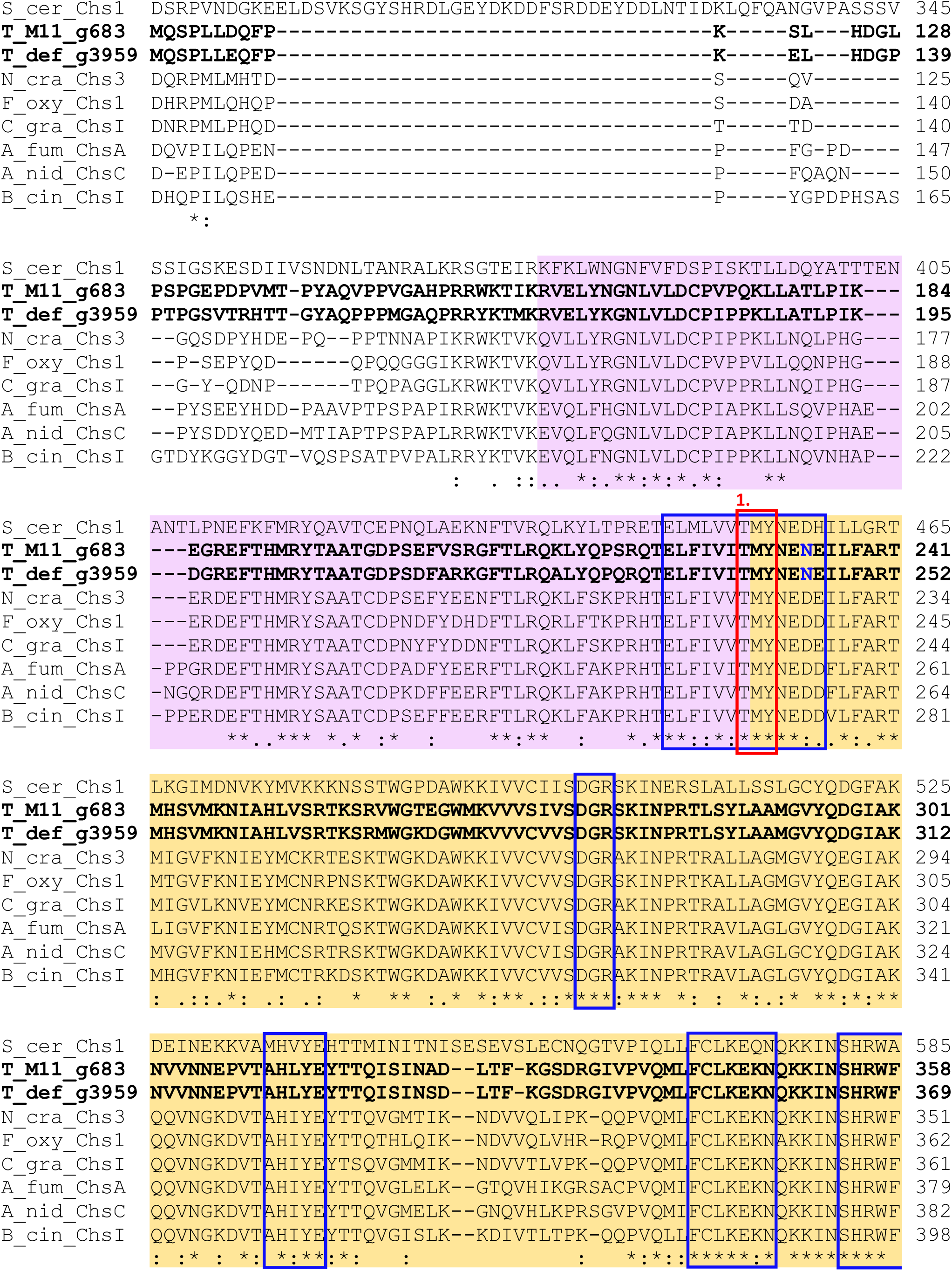
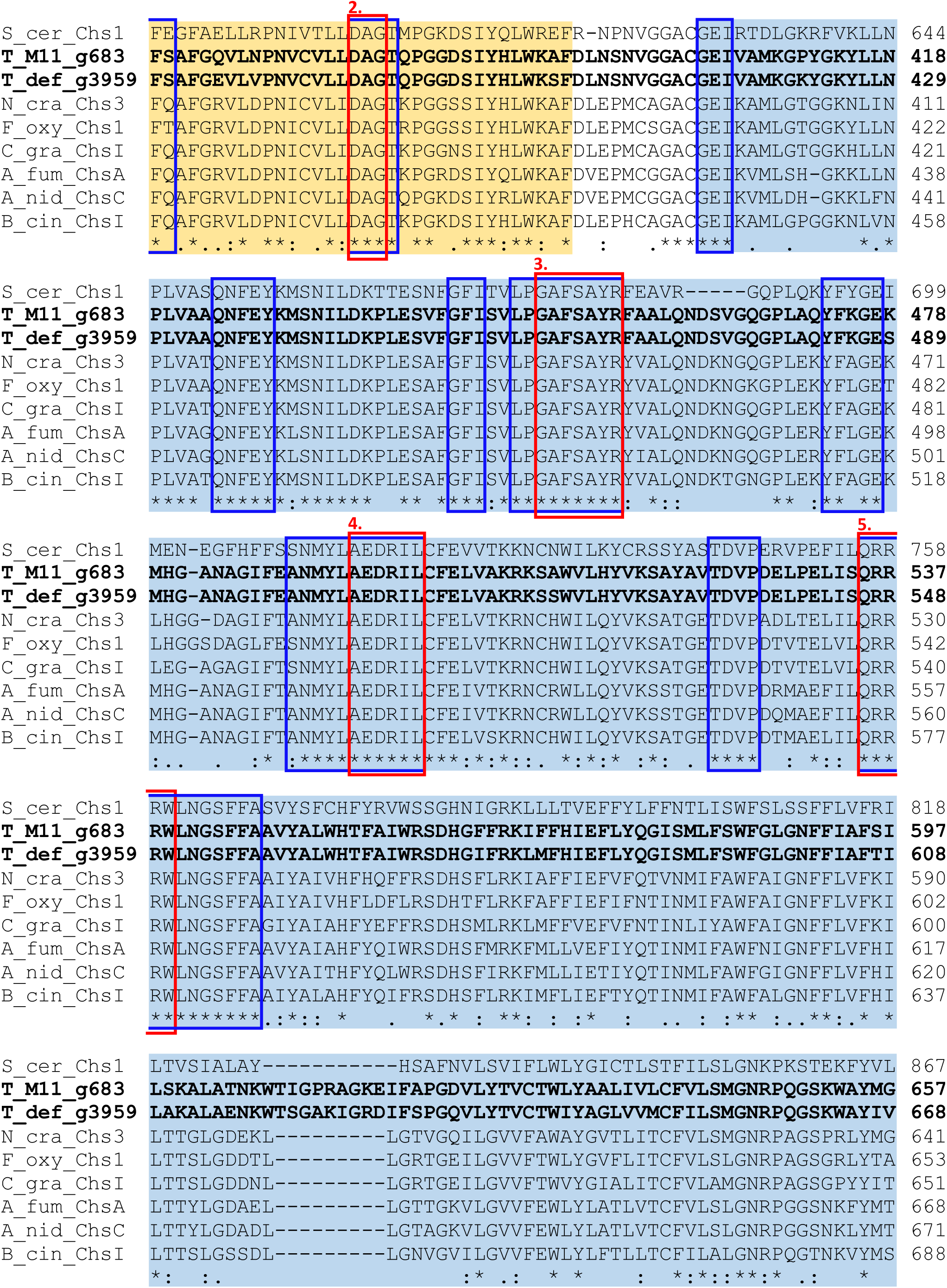
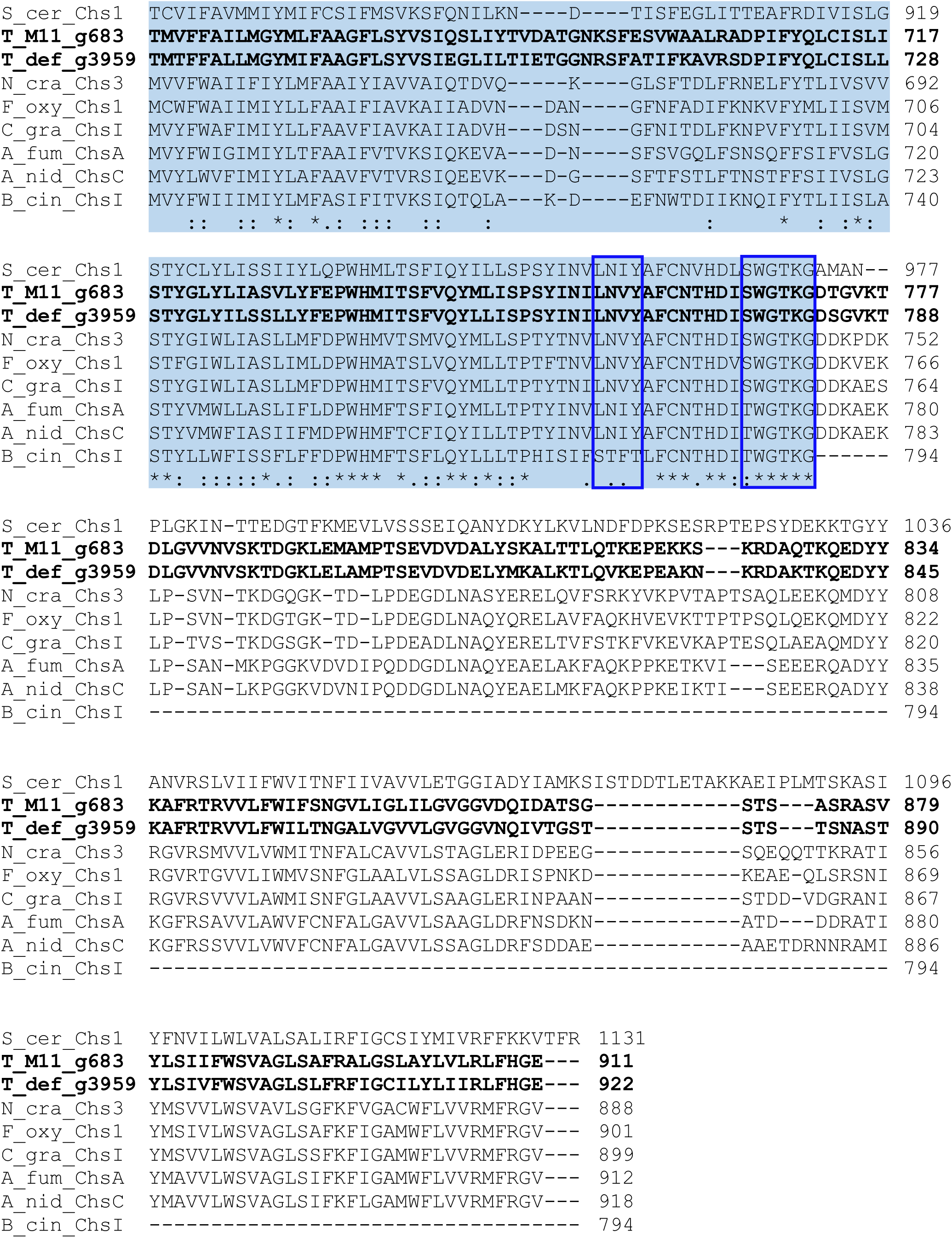
Sequence conservation in class I CHSs from *Taphrina* species. Multiple sequence alignment was performed with Clusal Omega. Conserved protein domains from the pfam database are higlighted as follows: pink, chitin synthase 1 N-terminal (pfam08407); yellow, chitin synthase1 (pfam01644); blue, partial chitin synthase2 (pfam03142). The start and end point of the domains pfam08407 and pfam01644 are from the NCBI database. The coordinates of chitin synthase2 are from Li et al. (2016), as the coordinates from the NCBI database were supported by alignment of only seven sequences. Red boxes indicate conserved functional motifs listed in Li et al. (2016): 1, ligand binding; 2, metal ion binding site; 3, donor saccharide binding; 4, acceptor saccharide binding; 5, product binding. Blue boxes indicate conserved sequence patterns defined by Li et al., 2016. Blue coloured amino acids in CHS sequences from *Taphrina* spp. do not match the conserved sequence patterns. Species abbreviations used: TM11, *Taphrina* species strain M11; Tdef, *Taphrina deformans*, Bcin, *Botrytis cinerea*; Foxy, *Fusarium oxysporum*; Ncra, *Neurospora crassa*, Cgra, *Colletotrichum graminicola*; Afum, *Aspegillus fumigatus*; Anid, *Aspegillus nidulans*, Scer, *Saccharomyces cervisiae*.

**Figure S4.**
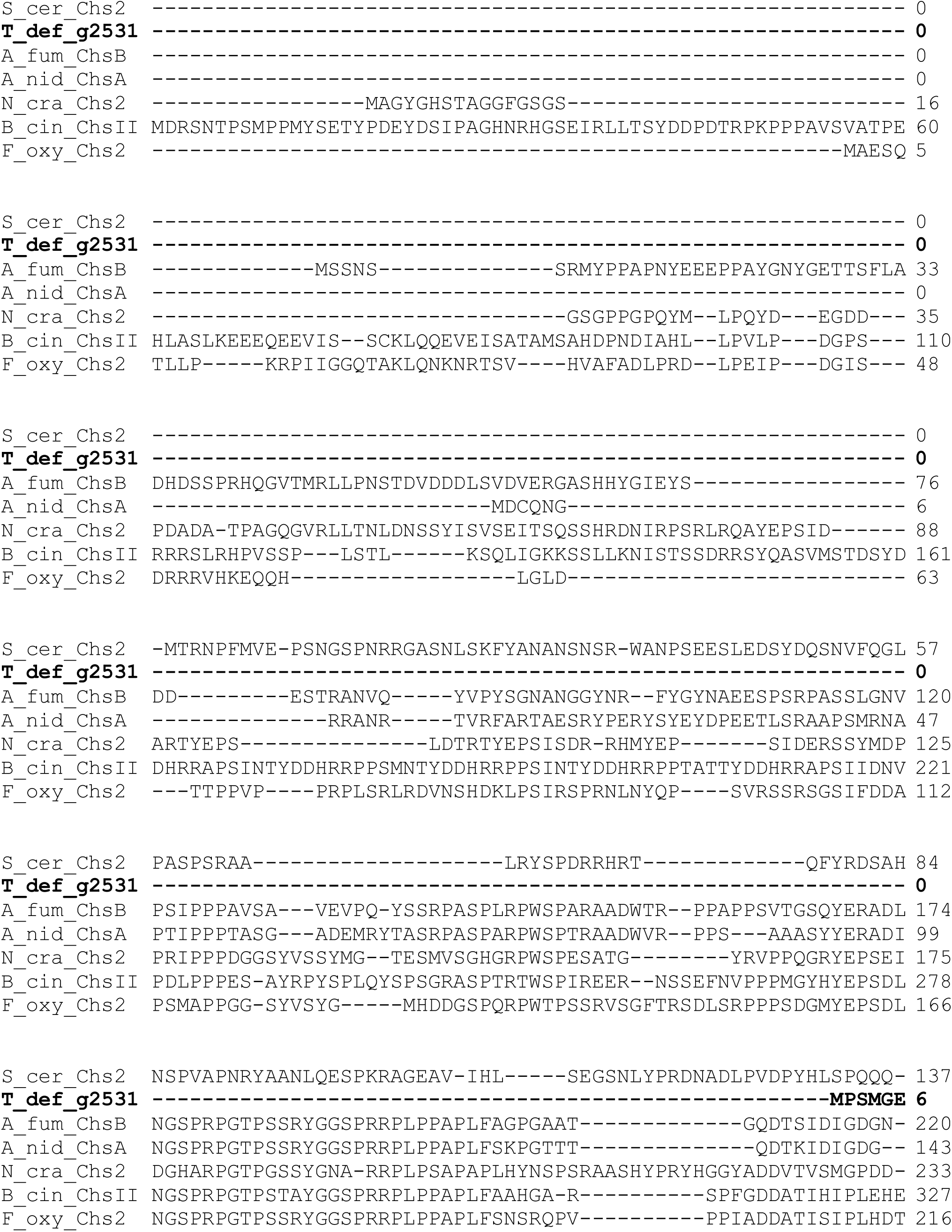
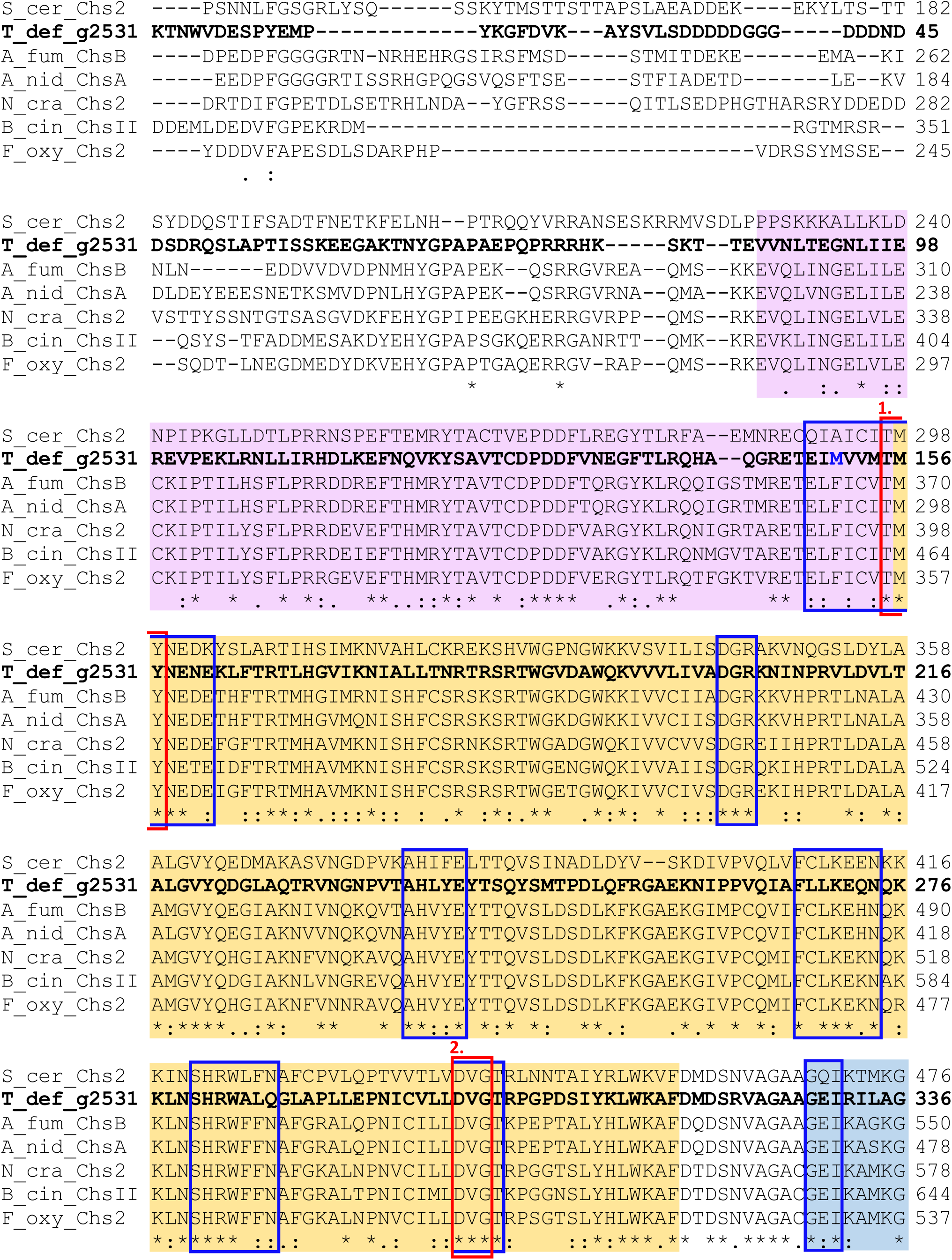
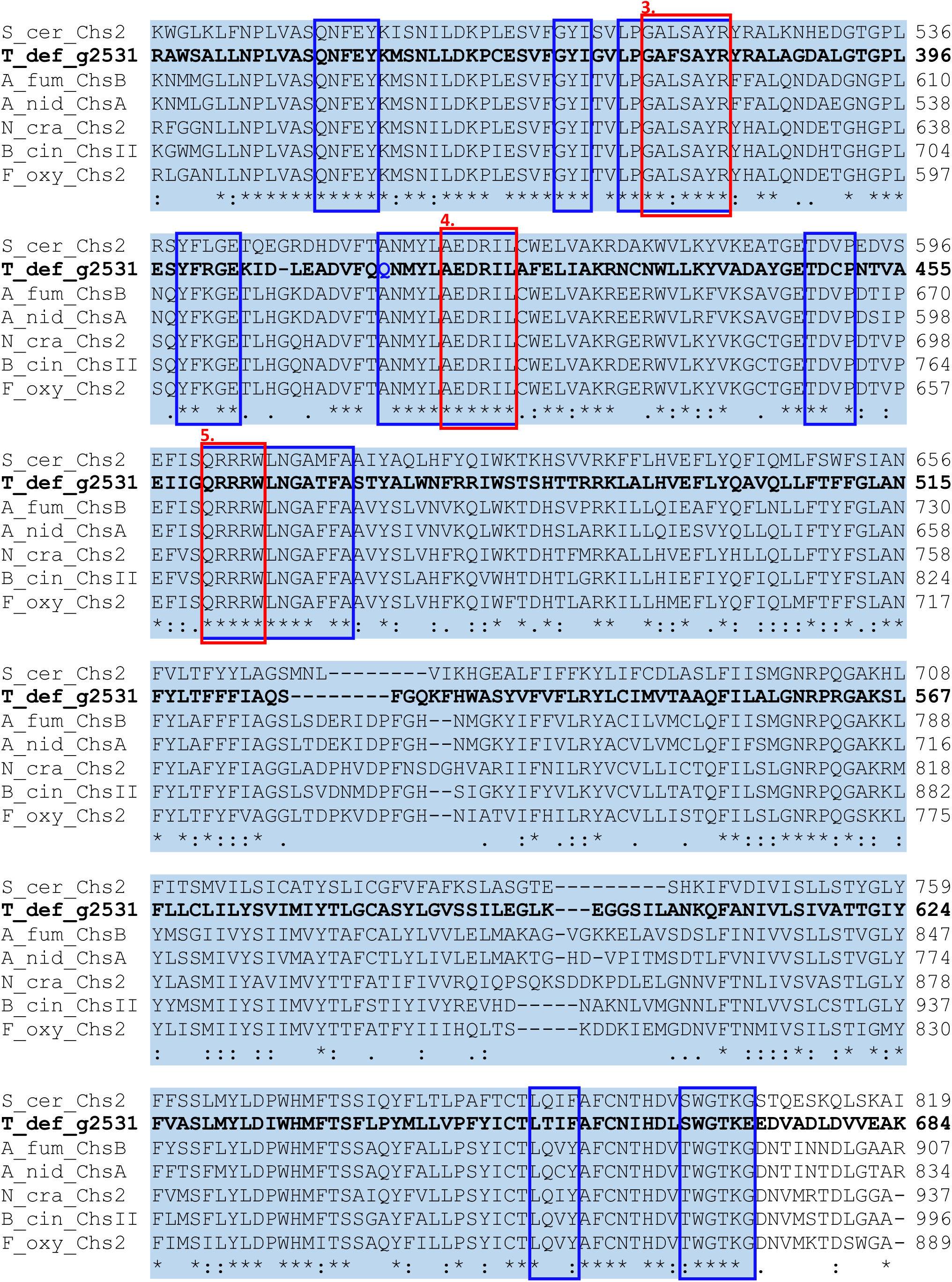
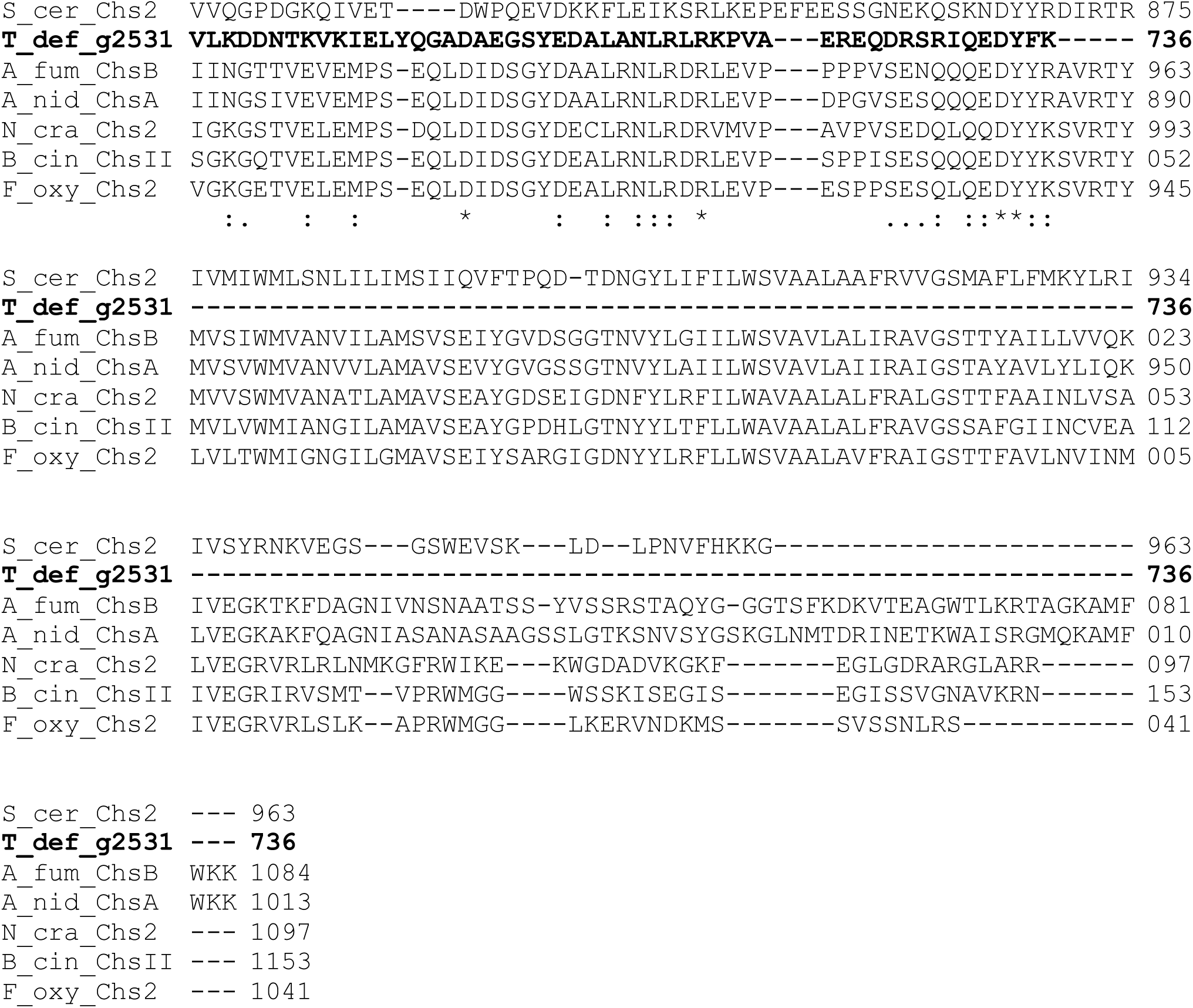
Sequence conservation in class II CHSs from *Taphrina deformans*. Multiple sequence alignment was performed with Clusal Omega. Conserved protein domains from the Pfam database are higlighted as follows: pink, chitin synthase 1 N-terminal (pfam08407); yellow, chitin synthase 1 (pfam01644); blue, partial chitin synthase 2 (pfam03142). The start and end point of the domains pfam08407 and pfam01644 are from the NCBI database. The coordinates of chitin synthase 2 are from Li et al. (2016), as the coordinates from the NCBI database were supported by alignment of only seven sequences. Red boxes indicate conserved functional motifs listed in Li et al. (2016): 1, ligand binding; 2, metal ion binding site; 3, donor saccharide binding; 4, acceptor saccharide binding; 5, product binding. Blue boxes indicate conserved sequence patterns defined by Li et al. 2016. Blue coloured amino acids in the CHS sequence from *Taphrina deformans* do not match the conserved sequence patterns. Species abbreviations used: Tdef, *Taphrina deformans*, Bcin, *Botrytis cinerea*; Foxy, *Fusarium oxysporum*; Ncra, *Neurospora crassa*, Afum, *Aspegillus fumigatus*; Anid, *Aspegillus nidulans*, Scer, *Saccharomyces cervisiae*.

**Figure S5.**
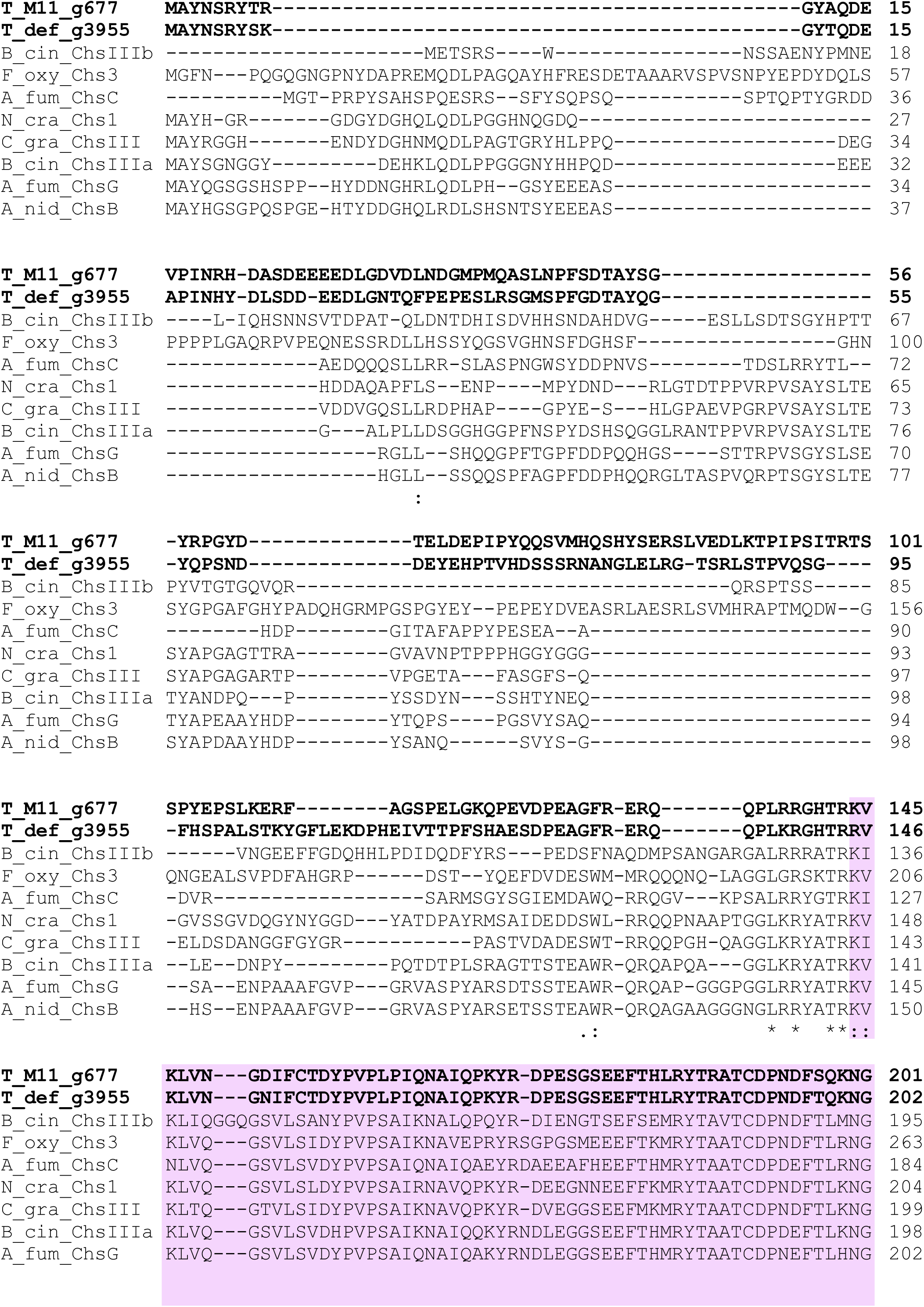
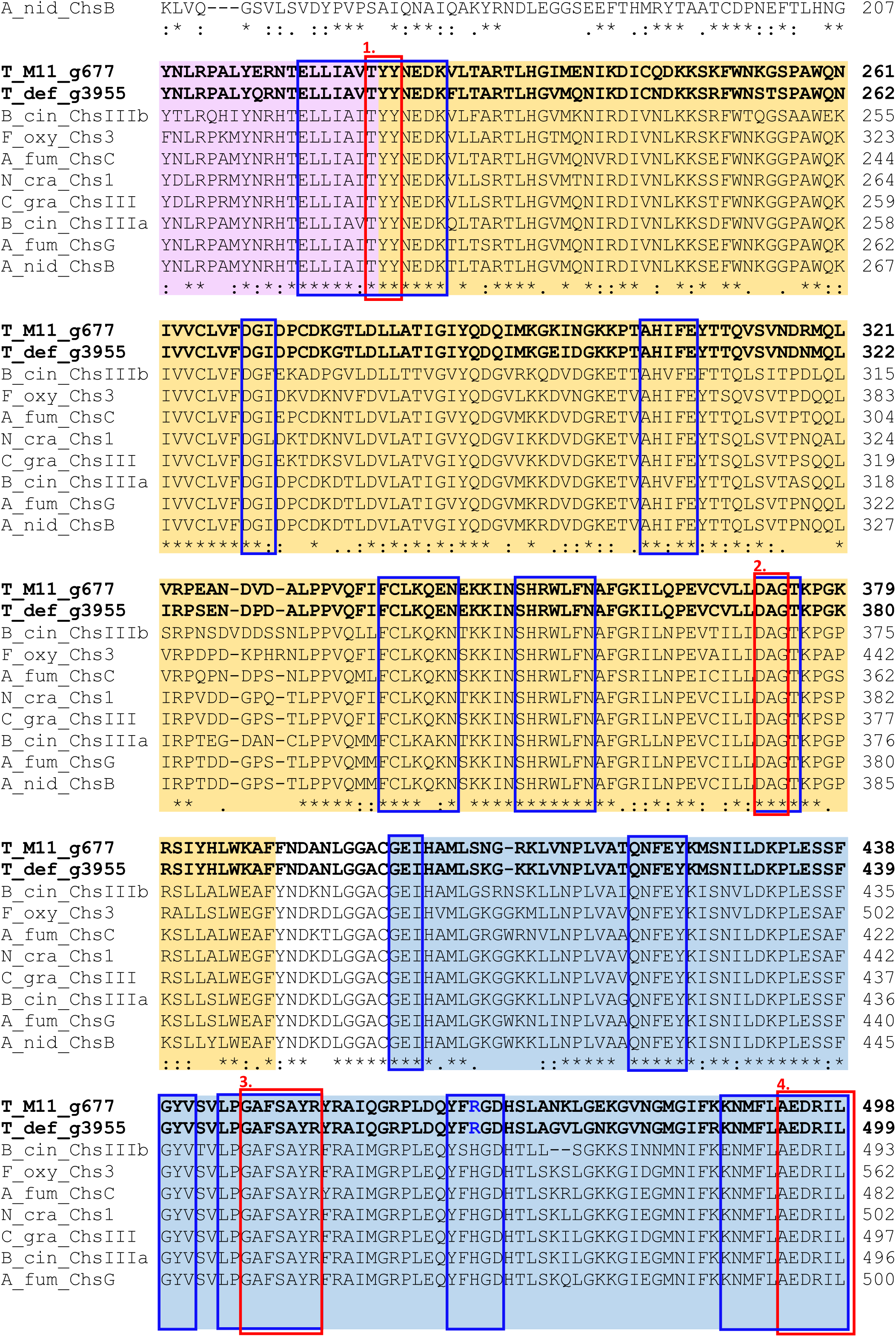
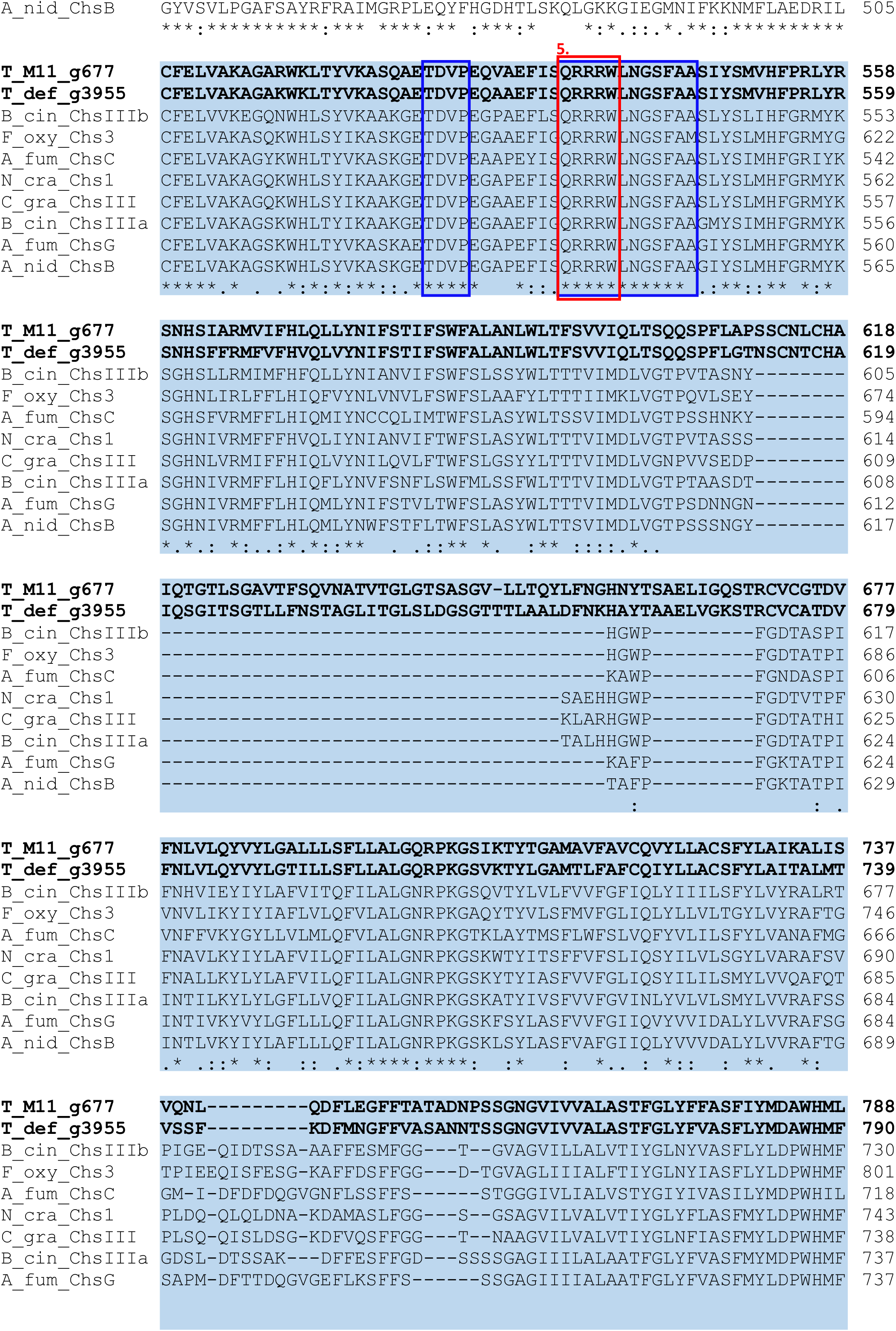
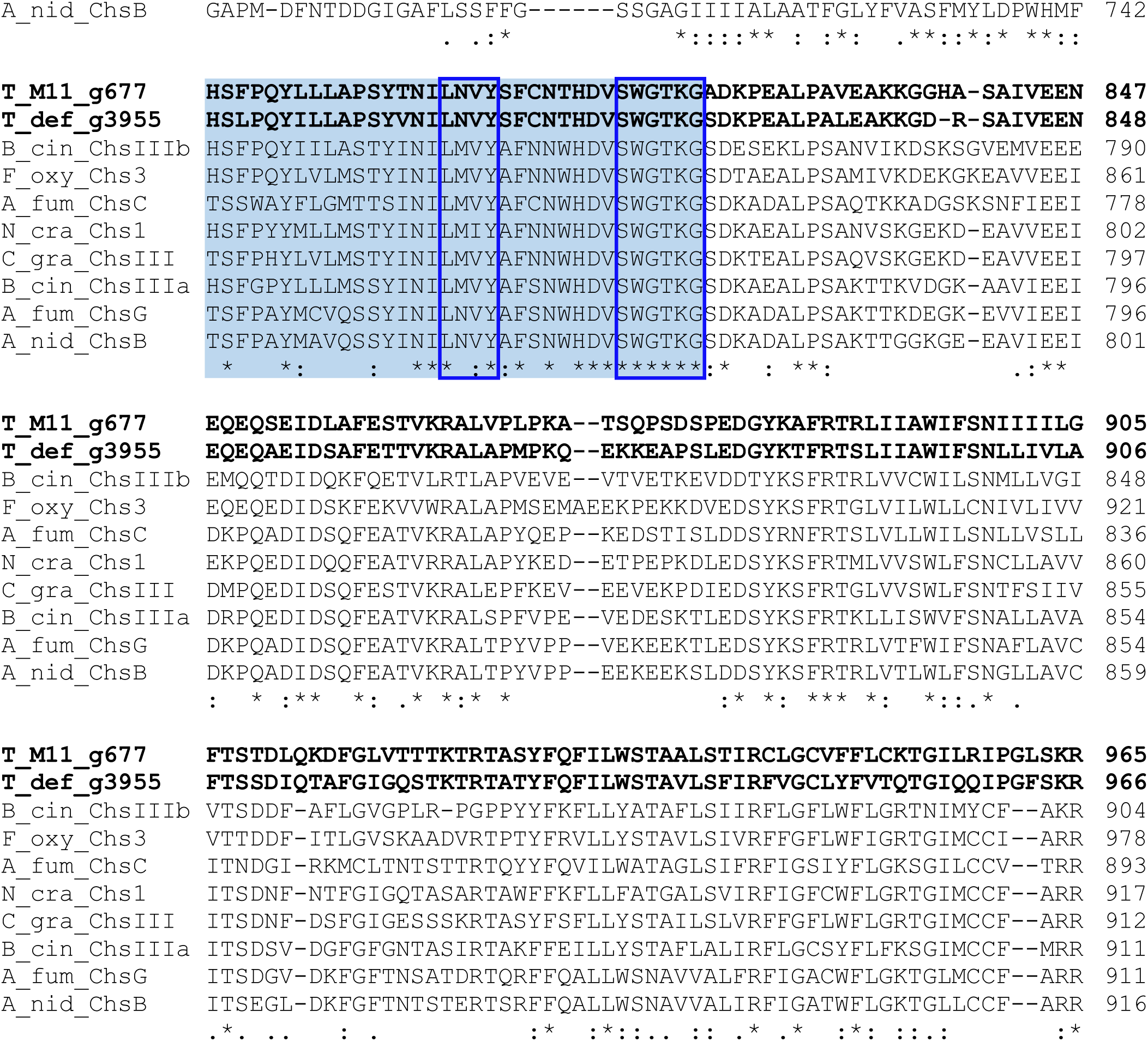
Sequence conservation in class III CHSs from *Taphrina* species. Multiple sequence alignment was performed with Clusal Omega. Conserved protein domains from the Pfam database are higlighted as follows: pink, chitin synthase 1 N-terminal (pfam08407); yellow, chitin synthase 1 (pfam01644); blue, partial chitin synthase 2 (pfam03142). The start and end point of the domains pfam08407 and pfam01644 are from the NCBI database. The coordinates of chitin synthase 2 are from Li et al. (2016), as the coordinates from the NCBI database were supported by alignment of only seven sequences. Red boxes indicate conserved functional motifs listed in Li et al. (2016): 1, ligand binding; 2, metal ion binding site; 3, donor saccharide binding; 4, acceptor saccharide binding; 5, product binding. Blue boxes indicate conserved sequence patterns defined by Li et al. 2016. Blue coloured amino acids in the CHS sequence from *Taphrina* species do not match the conserved sequence patterns. Species abbreviations used: TM11, *Taphrina* species strain M11; Tdef, *Taphrina deformans*, Bcin, *Botrytis cinerea*; Foxy, *Fusarium oxysporum*; Ncra, *Neurospora crassa*, Cgra, *Colletotrichum graminicola*; Afum, *Aspegillus fumigatus*; Anid, *Aspegillus nidulans*.

